# Mitochondria morphology changes dynamically during germ cell differentiation and under chemical stress in *C. elegans*: a quantitative ultrastructural study

**DOI:** 10.64898/2026.11.18.767639

**Authors:** Mathieu Vigneau, Vanessa Soldan, Arnaud Debernardi, Micol Guidi, Gaëtan Severac, Stéphanie Balor, Jean-Philippe Pradère, Anna Mattout

## Abstract

Mitochondrial dynamics are essential for cellular health and fertility, yet how mitochondrial ultrastructure adapts during gametogenesis and acute environmental stress remains poorly understood. Here, using high-pressure freezing electron microscopy and automated quantitative morphometry (*EMito-Metrix*), we mapped mitochondrial ultrastructure across the *Caenorhabditis elegans* germline and somatic tissues. We reveal that mitochondria undergo progressive structural maturation during germ cell differentiation, transitioning from small, round profiles to elongated networks with aligned cristae. Following short oxidative stress, differentiating germ cells and early embryos, unlike somatic or sperm cells, undergo a remarkable morphological remodeling to form protective, fusion-dependent donut-shaped and vacuolated spheroid mitochondria. Genetic disruption of the mitochondrial fusion factor *fzo-1* blocks this pathway and results in a collapse of stress-induced brood size. These results demonstrate that atypical mitochondrial morphotypes are not passive markers of damage, but an active, *FZO-1*-dependent defense strategy shielding female gametes. This work highlights the *C. elegans* gonad as a versatile model to explore mitochondrial plasticity, reproductive aging, and environmental toxicology.

## INTRODUCTION

Mitochondria are double-membrane organelles essential for cellular bioenergetics, signal transduction, calcium homeostasis, stemness, cell fate decisions, and apoptosis^1–3^. The inner mitochondrial membrane forms specialized invaginations termed cristae, which harbor the electron transport chain complexes and ATP synthase that drive oxidative phosphorylation^1–4^. Importantly, the fundamental mitochondrial features are deeply conserved among eukaryotes as well as the main enzymes that drive mitochondrial dynamics^4^. Mitochondrial dynamics, including fission and fusion which leads in general to small fragmented mitochondria or elongated tubular networks, are crucial for maintaining cellular health and function^4–6^. Specifically, while fusion alleviates damage by exchanging organellar contents across the network, fission facilitates quality control through targeted degradation (mitophagy) and supports cell division^4–9^. Mitochondrial dynamics seems particularly important in female germlines and fertility^10–13^. Throughout gametogenesis, across species, germ cells must undergo profound physiological transitions to support meiotic progression, differentiation, massive cellular growth, and the eventual energetic demands of fertilization and early embryogenesis^12^; but how mitochondrial architecture adapts to support these progressive changes remains poorly understood. Furthermore, under environmental stress or during reproductive aging, mitochondria in plants, *C. elegans,* mammalian and human oocytes frequently exhibit atypical morphologies, including the formation of vacuolated large spheroids or complex donut-shaped structures, also controlled at least partly by mitochondrial fusion enzymes^12,14–22^. However, a major biological question remains unresolved: are these atypical mitochondrial morphologies a symptom of age- and stress-induced dysfunction, or do they represent an active, conserved mechanism to protect fertility?

The 1 mm long famous nematode, *Caenorhabditis elegans* is a premier model organism for studying aging, stress responses and reproduction^23,24^. Particularly, because the *C. elegans* adult hermaphrodite gonad is spatially organized in a linear sequence of continuous development (**Figure 1A**), it offers an unparalleled *in vivo* system to study mitochondria features and dynamics during cellular differentiation, gametogenesis, and early embryogenesis^23,25^. Indeed, germ cell differentiation starts in the distal gonad from the mitotic distal tip, through meiotic prophase (pachytene and diplotene in the loop), to mature diakinesis oocytes in the proximal arm, before oocytes pass through the spermatheca to be fertilized and start the early embryonic development in the uterus (**Figure 1A**). Spermatogenesis with the resulting sperm cells stored in the spermathecae occurs in the fourth larval stage prior to oogenesis onset in the adult gonads. Recent advances have begun to shed light on the importance of mitochondrial remodeling through the gonad. For instance, it has been demonstrated that an initial wave of mitochondrial maturation, characterized by structural expansion and enhanced oxidative phosphorylation, is a critical driver of germline stem cell differentiation from the mitotic to the meiotic state in *C. elegans*^13^. However, while mitochondrial fluorescence used in that study^13^ and other studies, is extremely valuable for observing the network distribution, large-scale fragmentation dynamics in addition to other advantages, it lacks the resolution required to resolve internal cristae architecture or definitively identify complex morphological states. Consequently, two fundamental questions remain unanswered: First, how do mitochondria structurally remodel at the ultrastructural level during continuous gamete differentiation (from early germ cells to mature oocytes and early embryos) and compared to other cell types? Second, do differentiating germ cells structurally respond to environmental stress differently than terminally differentiated somatic cells?

**Figure 1:**
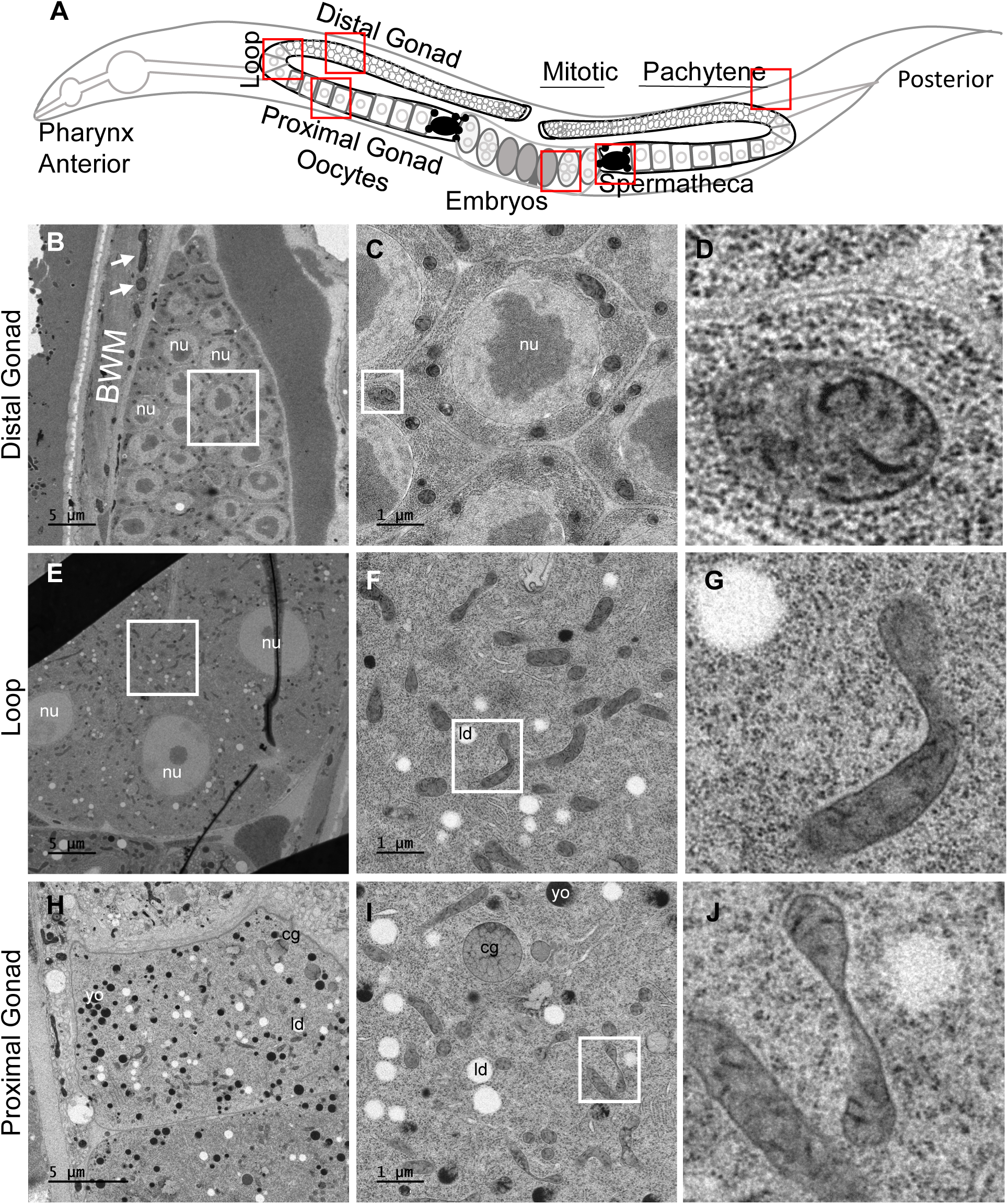
Mitochondrial morphology and ultrastructure in gonads of *C. elegans* at Adult Day 1 stage. A. Scheme of *C. elegans* hermaphrodite anterior and posterior gonads in early adulthood. Oocytes from both gonads pass through the spermatheca in order to self-fertilize and produce embryos that will start to develop *in utero*. B. Transmission electron micrograph of a longitudinal view of a part of the distal gonad. (nu) stands for nucleus; BWM for Bodywall Muscle, which are striated muscle under the cuticle that allow the locomotion of the worm. Arrows point to the mitochondria of the BWM cell. C. Enlargement of the region marked by a square in b. The numerous grey round organelles are mitochondria. D. Zoom in on one of the mitochondria from (C). E. Electron tomograph of the loop region of the gonad. The distal part is on the left and the proximal part on the right. F. Enlargement of the region marked by a square in e. The grey organelles are mitochondria. White droplets are lipid droplets (ld). G. Zoom in on one of the mitochondria from (F). H. Electron tomograph of longitudinal view of an oocyte in the proximal part of the gonad. (ld) are lipid droplets, (yo) are yolk, and (cg) stands for cortical granules. I. Transmission electron micrograph higher magnification of cytoplasmic content of an oocyte. Most of the grey organelles are mitochondria. (ld) are lipid droplets, (yo) are yolk, and (cg) stands for cortical granules. J. Zoom in on one of the mitochondria from (F). Mitochondria size from panels (C), (F), (I) are comparable given that those panels have the same scale bar.

To bridge this gap, we established an optimized, high-pressure freezing Transmission Electron Microscopy (TEM) procedure coupled with the use of EMito-Metrix, a high-performance automated pipeline for the segmentation and quantitative analysis of mitochondrial ultrastructure^26^. Using this approach, we mapped baseline mitochondrial morphometrics across the complete gonadal axis and in somatic body wall muscle. Notably, thanks to the gonadal linear organization and the small size of *C. elegans*, analysis within the exact same animals and samples can be achieved. We then challenged worms with brief, chemical oxidative stress (NaN3) prior to imaging to observe tissue-specific structural adaptations^27,28^. We provide a highly resolved quantitative map demonstrating that mitochondria undergo morphological and ultrastructural gradual changes, such as mitochondrial elongation and internal cristae shifts as they differentiate toward oocytes. Crucially, we demonstrate that while identical acute stress disrupts cristae organization globally across all tissues, only germ cells and early embryos undergo massive morphological remodeling to form among other atypical mitochondrial structures, ring or donut-shaped and vacuolated spheroid mitochondria previously associated with aging in both mammalian and *C. elegans* oocytes^12,14–19^. By assessing fusion-deficient *fzo-1* mutants, we reveal that this tissue-specific remodeling is an active, fusion-dependent defense mechanism necessary to preserve fertility under stress. This establishes the *C. elegans* gonad as a powerful model for defining the structural basis of female fertility and reproductive resilience.

## RESULTS

### Gradual morphological changes of mitochondria during differentiation of germ cells

In order to characterize mitochondria morphology and ultrastructure in different regions of the adult gonads of *C. elegans,* adult control hermaphrodite worms at day 1 of adulthood were observed with TEM, using several serial longitudinal sections of the animals from different blocs. High pressure freezing for cryo-fixation and substitution was applied to prepare the samples near native and physiological conditions and to obtain sufficient contrasted images to clearly detect fine morphology and the ultrastructure of mitochondrion.

We obtained highly resolved images (**Figure S1**). Given the small size of *C. elegans*, in addition to the gonads, the transmission electron micrographs allowed us to observe also other adjacent tissues and different cell types with a particularly fine resolution of different organelles such as the rough endoplasmic reticulum (**Figure S1A-C**); mitochondria (**Figure S1A-B**); nucleus and details such as heterochromatin patches juxtaposed to the nuclear periphery (**Figure S1D)** ; and examples of the inner and outer nuclear membranes with the latest decorated with 12 ribosomes in between two nuclear pores (**Figure S1E-F**). We could also capture abundant golgi apparatus in certain sections of certain cell types (**Figure S1G-H**); OP50 bacteria inside the lumen of the intestine of *C. elegans* (**Figure S1I**); pleated septate junctions (sj) in the spermatheca region (**Figure S1J**) as well as extracellular vesicles (EVs) trafficking in between sperm cells and the -1 oocyte^29,30^ (**Figure S1K-L**).

In the hermaphrodite *C. elegans* in order to characterize the mitochondria using this high resolutive TEM imaging during germ cell differentiation, we have focused our analysis in different gonadal regions where germ cells transition from a mitotic state in the distal region to differentiated states as they move proximally. Those regions include: the distal region of the gonad, the loop (or U turn) and the proximal region of the gonad where oocytes are formed in adult worms, as well as on mitochondria of sperm cells, in or around the spermatheca (**Figure 1A**). We have also focused our analysis on mitochondria of young embryos located in the uterus and of the well characterized mitochondria of the somatic body wall muscle cells, for comparison (**Figure 1A**).

Our observations indicate that mitochondria from the germ cells from either the distal, loop or proximal regions as well as from sperm cells are smaller than mitochondria from the body wall muscle (BWM) cells in all the different worms, gonads and sections we have observed (**Figures 1-3**). We could also find that when comparing mitochondria in the distal, loop and proximal regions -*i.e* comparing mitochondria during germ cells differentiation that led to oocyte formation-, mitochondria seem more homogeneous in shape and size in the distal part (**Figure 1B-D**) than in the loop and proximal part (**Figure 1E-J**), getting elongated with a tubular shape, or bended and twisted (or thinner in the middle) shapes (**Figure 1H-J**).

### Other organelles changes during germ cells differentiation

Other organelles exhibit changes during germ cell differentiation along the gonads. Lipid droplets (LDs) which are distinct cellular organelles consisting of a core of neutral lipids surrounded by a monolayer of phospholipids and used as fat storage^31^ are visible in germ cells mostly from the loop region and accumulate even more in oocytes in the proximal gonadal region and still found in cells of young embryos (**Figure 1 and Figure 2**).

**Figure 2:**
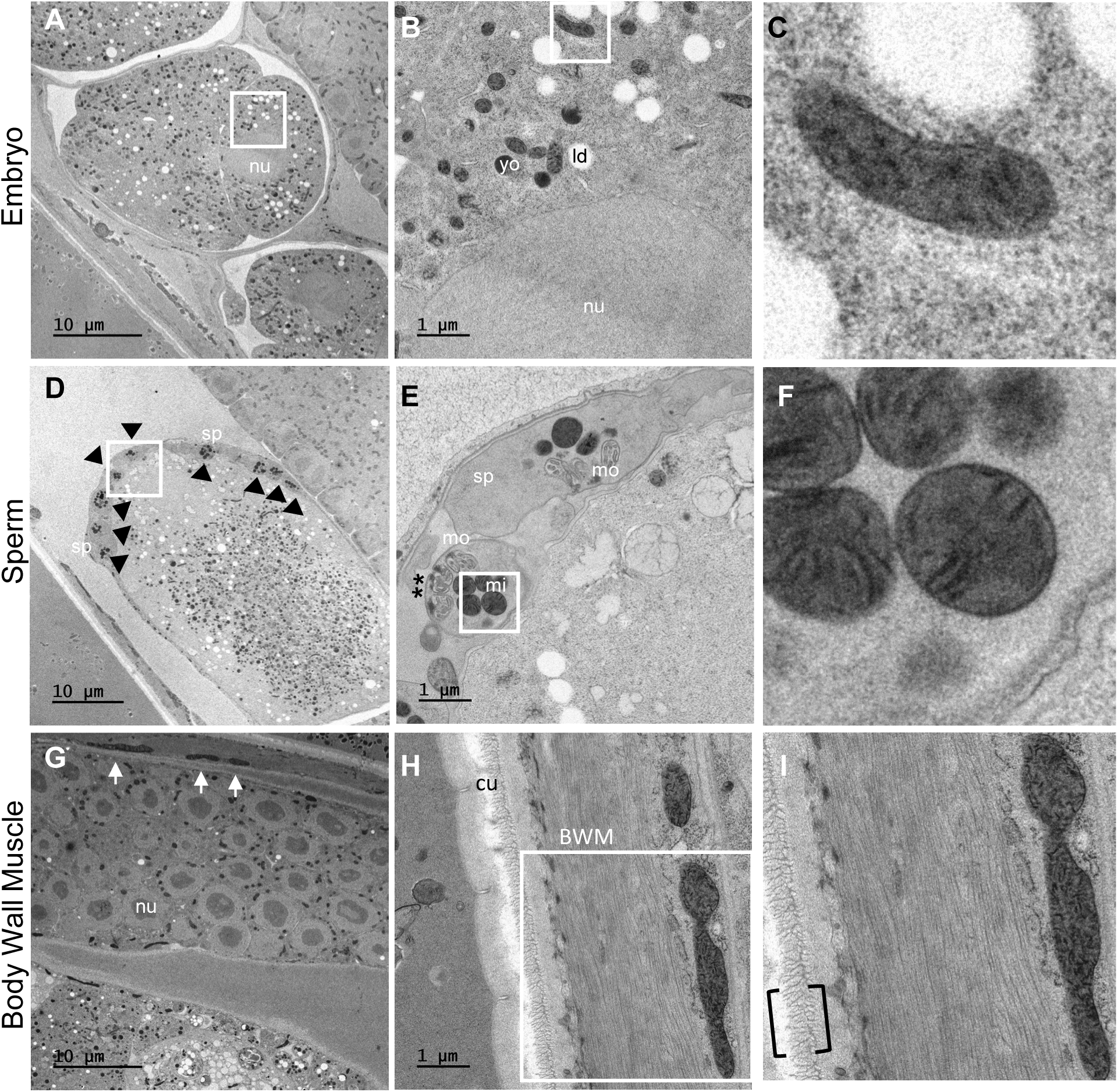
Mitochondrial morphology and ultrastructure in embryo, sperm and body wall muscle cells of *C. elegans*. A. Transmission electron micrograph (TEM) of a longitudinal view of a 4 cells-stage embryo. (nu) stands for nucleus. The region marked by a square is enlarged in b. The embryo is located in the uterus, next to the distal gonadal part (upper right corner). B. Enlargement of the region marked by a square in a. The grey organelles are mitochondria. White droplets are lipid droplets (ld) and darker droplets are yolk particles (yo), which are both abundant in the proximal gonad and embryos, especially compared to the distal gonad region as seen in (A). C. Zoom in of one of the mitochondria from (B). D. Transmission electron micrograph of a longitudinal view of the spermatheca. The spermatheca is inflated with the proximal oocyte passing through it. Spermatids (sp; black arrowheads) are observed in the spermatheca. The region marked by a square is enlarged in (E). E. Enlargement of the region marked by a square in (D). Two sperm cells are visible but their nuclei are not visible in that section of the bloc. The grey organelles are mitochondria (mi) and the white structured organelles are the membranous organelles (mo). Some of the collars (**) of the membranous organelles have fused to the spermatid plasma membrane, which occur during spermatid activation. F. Zoom in on one of the sperm mitochondria from (E). G. Electron tomograph of longitudinal view of a part of the body wall muscle (BMW) close to the distal gonad. Arrows point to mitochondria of the BWM, which are bigger and longer than mitochondria from the distal gonadal part found just beneath, observable few micrometers below. H. A higher magnification of body wall muscle fibers and of one mitochondrion, located under the cuticle (cu). I. Zoom in on one of the BWM mitochondria from h. The oriented collagen fibrils are probably what is observed in the brackets. Mitochondria size from panels (B), (E), (H) are comparable given that those panels have the same scale bar.

As in previous reports, we observed numerous yolk organelles (YOs) in oocytes and early embryos (**Figure 1 and Figure 2**), consistent with their role as nutrient/yolk-protein stores that support embryonic and early larval development^32,33^.

In *C. elegans*, cortical granules (CGs) are located at the cortex of maturing oocytes, as seen in (**Figure 1H-I**) and are specialized secretory vesicles that undergo synchronous exocytosis upon fertilization^34^. These granules deliver essential components, including chondroitin proteoglycans, to the zygote surface to facilitate the assembly of the eggshell, which acts as a barrier to prevent polyspermy and ensure embryonic osmotic integrity^35^.

### Mitochondrial morphological differences between germ, embryonic, sperm and muscle cells

Next, when comparing mitochondrial morphology in young embryos, sperm cells and body wall muscle cells **(Figure 1A**, **Figure2**) with those of the different parts of the gonads (**Figure 1B-J**) we could notice that mitochondria from embryonic cells (**Figure 2A-C**) are the most similar to the mitochondria in oocytes prior to fertilization in the proximal gonadal part (**Figure 1H-J**), consistent with the fact that in *C. elegans* as in mammals mitochondria are inherited from the female gamete only^36^. Some elongated mitochondria are still abundant in the embryonic cells **(Figure 2A)**, similarly to in oocytes **(Figure 1C)** compared to in pachytene germ cells in the distal gonad region (**Figure 1A**).

The hermaphrodite adult gonad contains both mature sperm and oocytes, enabling self-fertilization. Spermatogenesis occurs during the fourth larval stage (L4), and the resulting spermatids are stored in the spermatheca^37^. Upon the transition to adulthood, the gonad switches to oogenesis. In young and gravid adults, spermatids are stored in the spermatheca at the proximal-most end of the gonad (**Figure 1A**), where they activate to form mature and motile spermatozoa^38^. Mitochondria in sperm cells have a very distinct morphology compared to mitochondria in the female germ cells in the gonads. Mitochondria from sperm cells are very uniform in size and shape, with a remarkable round shape (**Figure 2D-F**). Sperm cells could be observed in the spermatheca as well as in proximity to the oocyte -1 and to the first(s) embryo(s) in all worms we could image the spermatheca and the oocyte and embryos surrounding it (**Figure S2A**), confirming that not all sperm cells are located in the spermatheca^39^. **Supplemental Figure 2B-C** shows an example of a sperm cell in close proximity to the first egg; the vitelline layer of the embryo eggshell appears to envelop this cell, potentially capturing the moment of fertilization and sperm entry.

Ultrastructural analysis reveals that *C. elegans* sperm cells are devoid of most canonical organelles, possessing primarily mitochondria and membranous organelles (MOs)^40^. Unlike typical cells, the spermatid cytoplasm lacks ribosomes and the endoplasmic reticulum (**Figure S2D-E**), aligning with the well-documented cessation of translation in mature sperm^37^. Another hallmark of *C. elegans* sperm, previously characterized^30,41^, is the absence of a nuclear envelope, that we can also perfectly capture with our high resolutive images (**Figure S2E**). This represents a remarkable exception to standard eukaryotic cellular organization. Rather than a membrane, the condensed nucleus of spermatids and spermatozoa in *C. elegans* is surrounded by a peripheral halo (**Figure S2E**).

Finally, mitochondria of somatic, body wall muscle cells were observed. Those cells, similarly to the oocytes require high amounts of energy in *C. elegans,* even if their relative requirement have not been assessed yet. Interestingly, those mitochondria that located next to the longitudinal striated fibrillar myofilaments of the muscle were much bigger and longer than any of the mitochondria in the other cells we have observed (**Figure 2G-I, Figure S3A**).

Altogether, mitochondria structurally mature during germ cell differentiation, culminating in elongated shapes within oocytes, while sperm and somatic cells retain distinct, tissue-specific mitochondrial architectures.

### EMito-Metrix analysis quantitatively supports increasing elongated mitochondria during germ cell differentiation

We recently developed [EMito-Metrix](https://www.emitometrix.org/)^26^, a high-performance and user-friendly automated pipeline that extracts quantitative data from 2D transmission electron microscopy (TEM) images. Previously used to compare and evidence muscular mitochondria differences – morphology and ultrastructure – across species^26^, EMito-Metrix represents a powerful tool for obtaining robust quantitative characterizations of mitochondria to complement our qualitative observations. After fine-tuning our deep learning model using a subset of our own TEM images, the pipeline performs automated segmentation for reliable mitochondrial detection. Once segmented, the software automatically calculates 24 distinct metrics per mitochondrion (**Table S1**) and provides a graphical interface for easy data visualization. An additional machine learning component utilizes predictive analytics to forecast how specific experimental conditions will affect mitometrics.

We applied this pipeline to analyze a large dataset of ∼2800 mitochondria across various cell types, averaging 475 mitochondria per cell type/tissue in control worms. Specifically, we analyzed 733, 504, 745, 558, 122, and 190 mitochondria in the distal gonad, loop region, proximal gonad, embryo, sperm, and muscle cells, respectively (**Figure S3**).

To ensure robust biological and technical sampling, images were acquired from two independent resin blocks prepared for TEM, encompassing three different adult worms and a total of four analyzed gonad arms. Imaging was performed across two to four distinct grids for each arm, separated by z-sections of at least 80 nm, though these additional acquisitions served primarily as a qualitative gauge of 3D mitochondrial shape and cellular context rather than for exhaustive quantification, with final analysis restricted to just one z-section per arm or two when well-separated in z, to reduce potential double-counting of individual mitochondria. All images used for quantitative analysis in the study were acquired using identical settings at 3,000x magnification, such as in **Figure 3A**.

**Figure 3:**
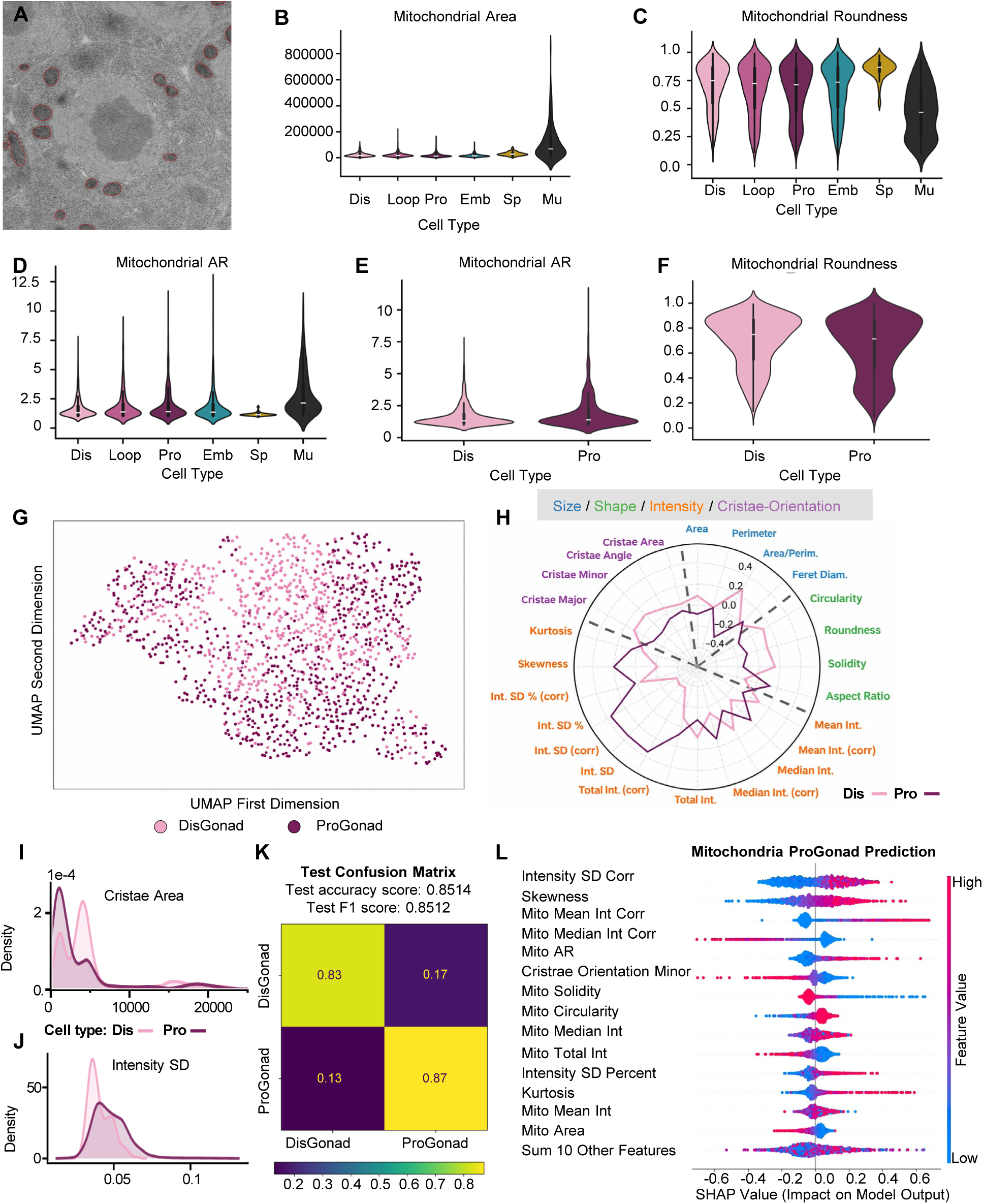
EMito-Metrix analysis show increasing elongated mitochondria and a change in cristae organization during germ cell differentiation. A. High magnification EM pictures of cells in the distal gonadal part early during germ cell differentiation. The image is overlaid with the automatically segmented and recognized mitochondria, delimited by a red line. Magnification 3000x. B. Mitochondria Area, a metric related to the size of the mitochondria is displayed as violin plots, for all the cell types (Dis: undifferentiated germ cells in the distal gonad; Loop: germ cells in the loop region of the gonad; Pro: oocytes in the proximal gonadal part; Emb: embryonic cells form early embryos in the uterus; Sp: sperm cells; Mu: body wall muscle cells ) in control conditions. C. Mitochondria Roundness related to the shape of the mitochondria are displayed as violin plots for all the cell types in control conditions. D. Mitochondria Aspect Ratio, also related to the shape of the mitochondria are displayed as violin plots for all the cell types in control conditions. E. Focused and enlarged view of the Aspect Ratio displayed as violin plots to compare mitochondria shape in germ cells early and late during germ cell differentiation, in the distal gonadal part and proximal gonadal part, respectively. F. Similar display that depicts the Roundness. G. UMAP projections of mitochondria metrics to compare each group. (733 mitochondria form the distal gonad, and 745 form the proximal gonadal part in oocytes.) UMAP embedding: Dimensionality reduction technique (UMAP) projecting all 24 quantitative mitochondrial descriptors per organelle into a 2D space, where spatial proximity reflects overall morphometric and ultrastructural similarity. H. Radar plot displaying all 24 mitometrics for both the germ cells early and late during germ cell differentiation, in the distal gonadal part and proximal gonadal part, respectively. Each spoke shows cell-type Dis and Pro means standardized to the common mean (0). The plot center represents the minimum plotted value minus a margin. The metrics can enter four categories, which are color-coded. Metrics describing the size are in blue, the shape in green, the Intensity / Texture/ Cristae density in orange, and the Cristae Orientation in purple. I. The Cristae Area metrics related to the Cristae organization of the mitochondria is displayed as density plots, to compare it in germ cells early and late during germ cell differentiation, in the distal gonadal part and proximal gonadal part, respectively. The x-axis represents the measurement in its unit (See Table S1 for all metrics units), and the y-axis (Density) represents the proportion of mitochondria per unit of x, with the total area under each curve equaling 100% of the group’s mitochondria. J. The Intensity SD metrics related to the Intensity category is displayed as density plots, to compare it in germ cells early and late during germ cell differentiation, in the distal gonadal part and proximal gonadal part, respectively. K. Confusion matrix generated by MLP (Multilayer Perceptron) algorithm to predict mitochondria from distal or proximal gonadal regions, less or more differentiated germ cells, respectively. L. Global explicability (using MLP algorithm) of the prediction model of the mitochondria from the more differentiated germ cells (Pro) compared to the (Dis) ranking the top-14 most important variables (displayed in order of importance based on the mean of absolute SHAP values)-. A positive or negative SHAP value on the x-axis means that the variable contributed to the positive or negative estimation of the group prediction, here the mitochondria of the oocytes in the proximal gonad. See Table S1 for metrics description and units.

The 24 current metrics measurable per mitochondrion using EMito-Metrix span four categories: size, shape, intensity/texture, and cristae organization (**Table S1** and **Figure S3**). Comparing all cell types analyzed in control animals, the data reveal a striking difference in mitochondrial size and area measured in 2D: muscle cell mitochondria are markedly larger than those found in any region of the gonad, oocytes, sperm, or embryonic cells (**Figure 3B** and **Figure S3**). This distinction is further supported by elevated perimeter, and Feret diameter, representing maximum mitochondrial length, in muscle compared to all other conditions (**Figure S3**). A clear variation is also evident in the roundness parameter, with values closer to 0.0 in muscles, indicating more elongated mitochondria. Sperm mitochondria exhibit roundness with the values the closest to 1.0 among all types analyzed (**Figure 3C**), consistent with qualitative observations (**Figure 2**). The circularity, which is a closely related measure to roundness, also clearly points to highly homogeneous circular mitochondria in sperm cells compared to in other cell types (**Figure S3**).

Beyond the uniform round shape of sperm mitochondria, violin plots measuring the roundness on the distal, loop, and proximal regions of the gonad show increasingly inflated lower tails (**Figure 3C**). This demonstrates that an increasing population of elongated mitochondria emerges during germ cell differentiation from pachytene through oogenesis, even though the majority retain a roundish morphology in 2D TEM images. These quantitative results corroborate the qualitative findings described in **Figures 1** and **2**. Analyzing the aspect ratio (AR) that represents the length-to-width ratio, where a high AR indicates tubular/filamentous mitochondria consolidates those conclusions (**Figure 3D**). Mirroring the roundness trends, the AR violin plots across the distal, loop, and proximal gonadal regions show an inverse elongation with increasingly inflated upper tails from the distal to the proximal gonad (**Figure 3D**). This captures again the progressive elongation and diversification of a subset of mitochondria during germ cell differentiation from the distal to the proximal gonad. A subset of elongated mitochondria is also detectable in embryonic cells of Day 1 adult worms following the fertilization of proximal oocytes, which typically occurs within minutes.

A focused one-on-one comparative analysis between the early and end points of the germ cell differentiation, with early meiotic cells in the distal gonad and oocytes in the proximal gonad highlights well the subset of mitochondria with the violin tails, that are the most elongated and the least circular/round in the differentiated germ cells (**Figure 3E-F**), while the median also shows similar trends with an AR value of 1.66 vs 1.89 in distal vs proximal gonadal mitochondria and a roundness of 0.69 vs 0.65. A UMAP dimensionality reduction graph alongside a radar plot displaying multi-parametric mitochondrial metrics (**Figure 3G** and **H**, respectively) clearly provide the distinct relative profiles of mitochondria between the two conditions. UMAP (Uniform Manifold Approximation and Projection), which is a dimensionality reduction technique which transforms the 24 metrics of each mitochondrion into a simpler 2D representation helps to identify distinct distribution of Pro- and Dis-Gonad populations, and confirms using an unsupervised analysis how both mitochondria populations display morphological divergence (**Figure 3G**). More specifically, using multi variate depiction with the radar plot, mitochondria in oocytes, in the proximal gonad exhibit higher values for the aspect ratio (AR), whereas circularity, roundness, and solidity are highest in the undifferentiated germ cells that are found in the distal gonad (**Figure 3H**), further supporting morphological differences between the distal and proximal gonadal parts.

### Cristae undergo progressive structural reorganization during germ cell differentiation

The ultrastructural changes in Crista’s orientation, alignment and density within the mitochondria is captured by the four Cristae Orientation measures and intensity measures (**Table S1; Figure 3H**). Interestingly, cristae metrics change and are on average moderately higher in the mitochondria of the distal region compared to the proximal gonad part (**Figure 3H**), suggesting a potential change in the ultrastructure organization. In addition, when the density plots of those cristae metrics are visualized, as exemplified in **Figure 3I**, the different organization of the cristae in the oocytes is captured. Thus, a swap in the prominent peaks of the density plot of the Cristae Area which represents the surface occupied by periodic cristae organization, stands for a change in the organization of cristae during germ cell differentiation (**Figure 3 I**).

Furthermore, to investigate cristae organization, the different metrics related to the intensity represent a proxy of the crista’s density within the mitochondria. Notably, the mean and the median intensity as well as the intensity standard deviation (Int SD) and their relative metrics increase in mitochondria during germ cell differentiation (**Figure 3H; Figure S3; Table S1**). For instance, mitochondria in the oocytes of the proximal gonad displays a higher overall intensity SD than the distal group, with a shift to the right (**Figure 3J**), suggesting that cristae density is increased in the proximal gonads compared to distal gonads. When looking at the peak width (**Figure 3J**), it demonstrates a broader distribution, with a broader peak in the proximal group, supporting a higher heterogeneity of cristae density repertoire in mitochondria populations. Given the high energy demand for oocyte and fertilization that occur shortly after, we propose that our observed quantitative changes in the fine structural and density reorganization of the cristae during germ cell differentiation is reflecting a higher mitochondria respiration^42^. This type of cristae organization would certainly contribute to the increase in ATP production, electrochemical potential as well as in ROS production recently shown in the proximal gonad compared to the distal gonad^13^.

Given that mitochondria of Distal and Proximal parts are relatively well segregated on the UMAP plot (**Figure 3G**), we have used the analytic prediction tool to test if a trained neural network can predict the cell type from which each mitochondrion originates, based on the metrics. The confusion matrix generated by MLP algorithm can predict the two categories with a high recall score/success rate of 83% and 87%, respectively after a training phase (**Figure 3K**), further confirming the clear differences of mitochondria morphometrics between the germ cells at different differentiation state in the Dis and Pro-Gonad. The SHAP (SHapley Additive exPlanations) values generated with EMito-Metrix gives the explicability of the prediction of the mitochondria in the proximal gonad, that classify in order of importance the metrics that contribute to that prediction. A positive or negative SHAP value on the x-axis means that the variable contributed to the positive or negative estimation of the condition prediction (**Figure 3L**). The Intensity related metrics (including Intensity SD and Mean or Median Int) as well as the Mito AR and Mito solidity are among the most predictive features, consistently with the metrics observed in the radar plot (**Figure 3H**) and to the observations of the elongated and bended shape that the mitochondria undertake during germ cell differentiation (**Figure 1**, **Figure 3D-E**).

Altogether, our TEM observations and quantitative analysis reveal distinct differences in mitochondrial morphology and ultrastructure across various *C. elegans* cell types, highlighted by larger muscular mitochondria and rounder sperm mitochondria compared to other analyzed populations. Importantly, these data demonstrate progressive changes during germ cell differentiation from early pachytene to later oogenesis stages, with a subset of mitochondria becoming elongated and bent as germ nuclei mature in day 1 adult animals, and a reorganization of the cristae.

### Stress drives tissue-specific mitochondrial remodeling to form donut and spheroid organelles in *C. elegans* germ cells

This striking mitochondrial remodeling within the gonads of adult worms was a fortunate discovery. Initially, we aimed to characterize *C. elegans* gonadal arms, seeking to capture longitudinal sections containing the distal, loop, and proximal regions in the same plane. To achieve flatter, straightened samples in our blocks and optimize longitudinal sectioning for cryo-fixation, we tested the use of 30 mM NaN_3_ for 15 to 45 minutes in cold M9 buffer to anesthetize the worms prior to cryo-fixation, supplementing our standard control condition of collection in cold M9 buffer alone. However, this mild concentration of NaN3 also inhibits mitochondrial cytochrome c oxidase (Complex IV) resulting in rapid ATP depletion and an altered cellular redox balance as recently demonstrated^28^. Importantly, this is not a lethal dose, and worms can fully recover from paralysis once the exposure is removed^28^.

Generally, internal tissue and cellular organization appear relatively normal in the same cell-types / tissues (**Figure 1A**) we have looked at in control conditions, or at least not drastically disrupted in the low-magnification overview panels (left columns, **Figures 4 and 5**). However, when examining mitochondrial morphology at higher magnification, we observed elongated and bent mitochondrial shapes as early as the distal gonadal region, similar to those detected later during normal germ cell differentiation in control young adult worms (**Figure 1E-J and Figure 4A-C**). In addition to these elongated structures, a wide array of atypical mitochondrial morphologies was observed throughout the distal, loop, and proximal regions of the gonad (**Figure 4, Figure S4 and S5**). These include dumbbell-shaped, C-shaped, cup-shaped, hooked, lasso-shaped or horseshoe-shaped (**Figure S5A**), and branched or interconnected mitochondrial networks, as well as more striking donut-shaped or ring-shaped structures that encompass cytoplasm containing ribosomes and occasionally other organelles (**Figure S5B**). Some of these donut-shaped profiles likely correspond to cross-sections of three-dimensional (3D) mitochondrial spheroids, with an internal vacuole or lumen, a conclusion supported by examples in **Figure S5C** and our analysis of serial Z-sections (spaced 80 nm apart), which reveal a varying lumen area across planes. Based on our observations, intermediate or partial bending states, such as dumbbell, cup, or half-ring shapes, frequently appear following elongation and prior to complete donut closure (**Figure S4 and S5A**). These findings are concordant with recent observations in other cell -types and organisms^43^. The formation of donut-shaped mitochondria in old *C. elegans* germ cells was shown to be dependent of mitochondrial fusion and of FZO-1 enzyme^19^.

**Figure 4:**
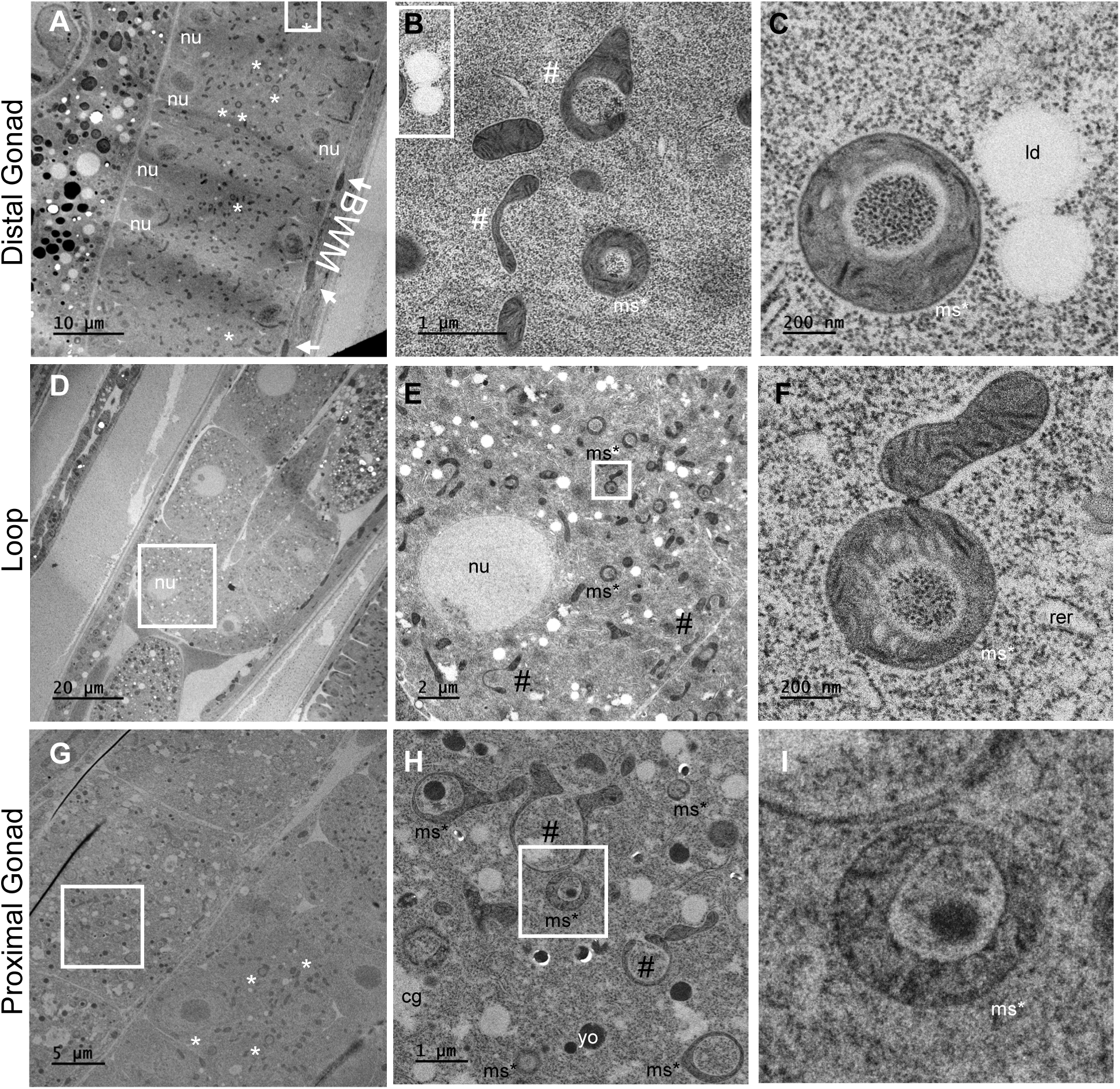
Mitochondrial morphology and ultrastructure in gonads of anesthetized *C. elegans* at Adult Day 1. A. Transmission electron micrograph (TEM) of a longitudinal view of a part of the distal gonad. (nu) stands for nucleus; BWM for Body wall Muscle. Arrows point to the mitochondria of the BWM cell. Asterisks (*) show examples of spheroid or donut shaped mitochondria visible at this magnification. B. Enlargement of the region marked by the white square in the upper part of panel a. The elongated mitochondria (#) as well as the spheroid or donut shaped mitochondria (ms*) are more clearly detectable at this magnification. C. Zoom in on one of the mitochondria just outside of panel b (white box). Two lipid droplets one next to each other are found in proximity to the spheroid mitochondria (ms*). This spheroid shaped mitochondria (ms*) seems to have embedded in its internal part some ribosomes from the cytoplasm. D. Electron micrograph of the loop region of the gonad. E. Enlargement of the region marked by a square in (D). (nu) : nucleus; (#) elongated, horseshoe or dumbbell shaped mitochondria examples; (ms*): spheroid / donut shaped mitochondria. F. Zoom in on mitochondria from the white boxed region in panel (E). G. TEM view of an oocyte in the proximal part of the gonad. Asterisks (*) show examples of spheroid or donut shaped mitochondria visible at this magnification. H. Higher magnification of the white boxed region of (G). (cg): cortical granules; (yo): yolk particles, some yolk proteins seem to be internalized in the hole of some spheroid/ donut shaped mitochondria (ms*), as we can see in two examples in that picture I. Zoom in on one of the mitochondria from (H). Cytoplasmic content seems to fill the interior part of the spheroid / donut shaped mitochondria (ms*).

**Figure 5:**
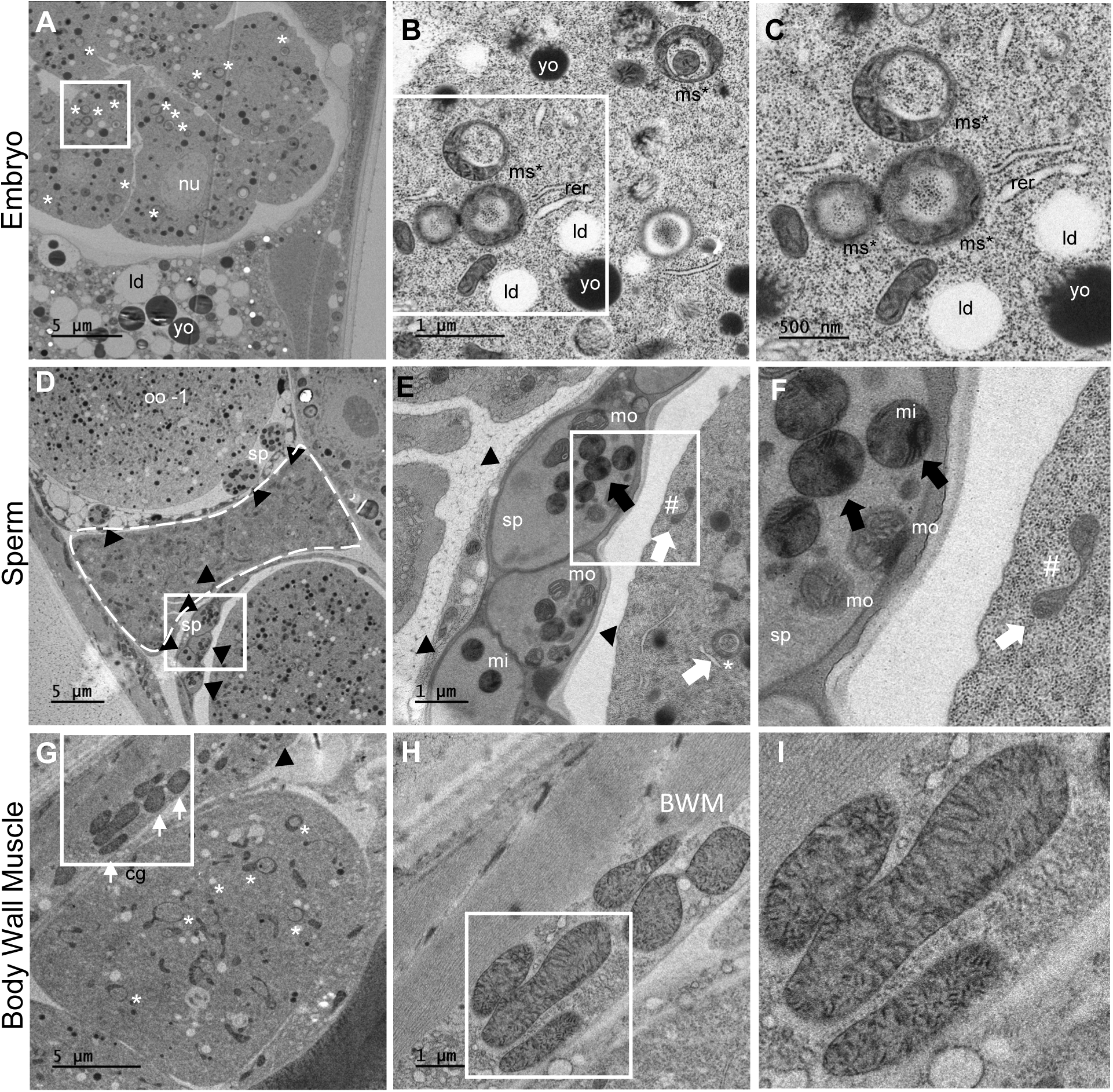
Mitochondrial morphology and ultrastructure in embryo, sperm and body wall muscle cells of anesthetized *C. elegans*. A. Transmission electron micrograph (TEM) of a longitudinal view of a young embryo. (nu) stands for nucleus. The region marked by a square is enlarged in b. The embryo is located in the uterus. Asterisks (*) show example of spheroid / donut shaped mitochondria visible at this magnification. B. Enlargement of the region marked by a square in a. The grey organelles are mitochondria. White droplets are lipid droplets (ld) and darker droplets are yolk particles (yo), which are both abundant in the proximal gonad and embryos. (rer): rough endoplasmic reticulum; (ms*): spheroid / donut shaped mitochondria examples. C. Zoom in mitochondria from b. Classical oval shaped mitochondria are detectable in addition to spheroid / donut shaped mitochondria (ms*). D. Transmission electron micrograph of a longitudinal view of the spermatheca (dashed line). Spermatids (sp; black arrowheads) are observed in proximity to the most proximal oocyte (oo -1) and in between the exit of the spermatheca and the first fertilized egg on the bottom part. The region marked by a square is enlarged in (E). E. Enlargement of the region marked by a square in (D). Three spermatids (sp; black arrowheads) are visible but their nuclei are not visible in that section. The dark grey organelles are spermatid mitochondria (one black arrow example), the lighter grey organelles on the egg part are mitochondria with an elongated dumbbell shape (#) and a spheroid / donut shape (ms*) and are marked by white arrows. The white structured organelles in the spermatids cell body are the membranous organelles (mo). F. Zoom in on four spermatid mitochondria from (E) and the elongated dumbbell shaped mitochondria (#) detectable few micrometers away from the round mitochondria of the spermatids. No ribosomes (back dots) are detectable in the cytoplasm of the spermatids in contrast to the cytoplasm of the other cell types surrounding the spermatids (E, F). G. Micrograph of a longitudinal view of a part of the body wall muscle (BMW) close to an oocyte in the proximal gonad region. Arrows point to typical mitochondria of the BWM, which are bigger and longer than mitochondria from the oocyte observable just beneath, many of which have spheroid/ donut shapes (*). H. A higher magnification of body wall muscle fibers and of mitochondria from the white square selection, located under the cuticle. I. Zoom in BWM mitochondria from (H).

These atypical mitochondrial shapes, particularly the donut-shaped structures, were also abundantly detected in the cells of *in utero* embryos (**Figure 5A-C**). Importantly, atypical mitochondrial morphologies were never observed in sperm cells or somatic cells (such as body wall muscle cells) across all animals and samples subjected to the same mild and short chemical stress condition (**Figure 5D-I**). Examples of typical mitochondria in sperm cells alongside atypical mitochondria in nearby embryonic cells (separated by only a few micrometers) are shown in **Figure 5E-F**. Similarly, typical mitochondria in muscle cells can be seen directly juxtaposed to oocytes that are packed with atypical mitochondria just microns away (**Figure 5G-I**).

Together, these findings indicate that mitochondria from undifferentiated germ cells, oocytes, and embryonic cells undergo dynamic and specialized shape changes in response to short and mild chemical stress. In contrast, neither sperm cells nor the somatic cells observed exhibit drastic conformational changes of mitochondria as observed in the germline and early embryos. This suggests that the appearance of atypical-shaped-mitochondria only in the gonads upon chemical stress might be regulated by a specific pathway that does not intervene in somatic cells.

To quantify this phenomenon, we imaged 10 gonads from 8 worms across 3 different blocks at different Z planes. Donut-shaped mitochondria were observed to a similar extent across all 10 gonads. To evaluate the abundance of atypical mitochondria in all cell types and conditions, we focused specifically on the donut-shaped population. Using the total number of mitochondria detected and segmented per condition by the EMito-Metrix pipeline, we counted the number of donut-shaped mitochondria across all analyzed images and calculated their relative abundance (**Figure 6A**). Under NaN_3_-chemical stress, we found that approximately 9% of the mitochondria in the distal gonad germ cells adopt a ring or donut shape, 10% in proximal gonad oocytes, and 20% (or 200‰) in young embryos, the highest proportion. In contrast, donut-shaped mitochondria were virtually absent (0%) in control conditions, with the exception of a very minor residual frequency (0.2%) observed as a rare event in embryos. Mitochondria in the loop region of the gonad were not quantitatively analyzed due to an insufficient sample size for a representative evaluation.

**Figure 6:**
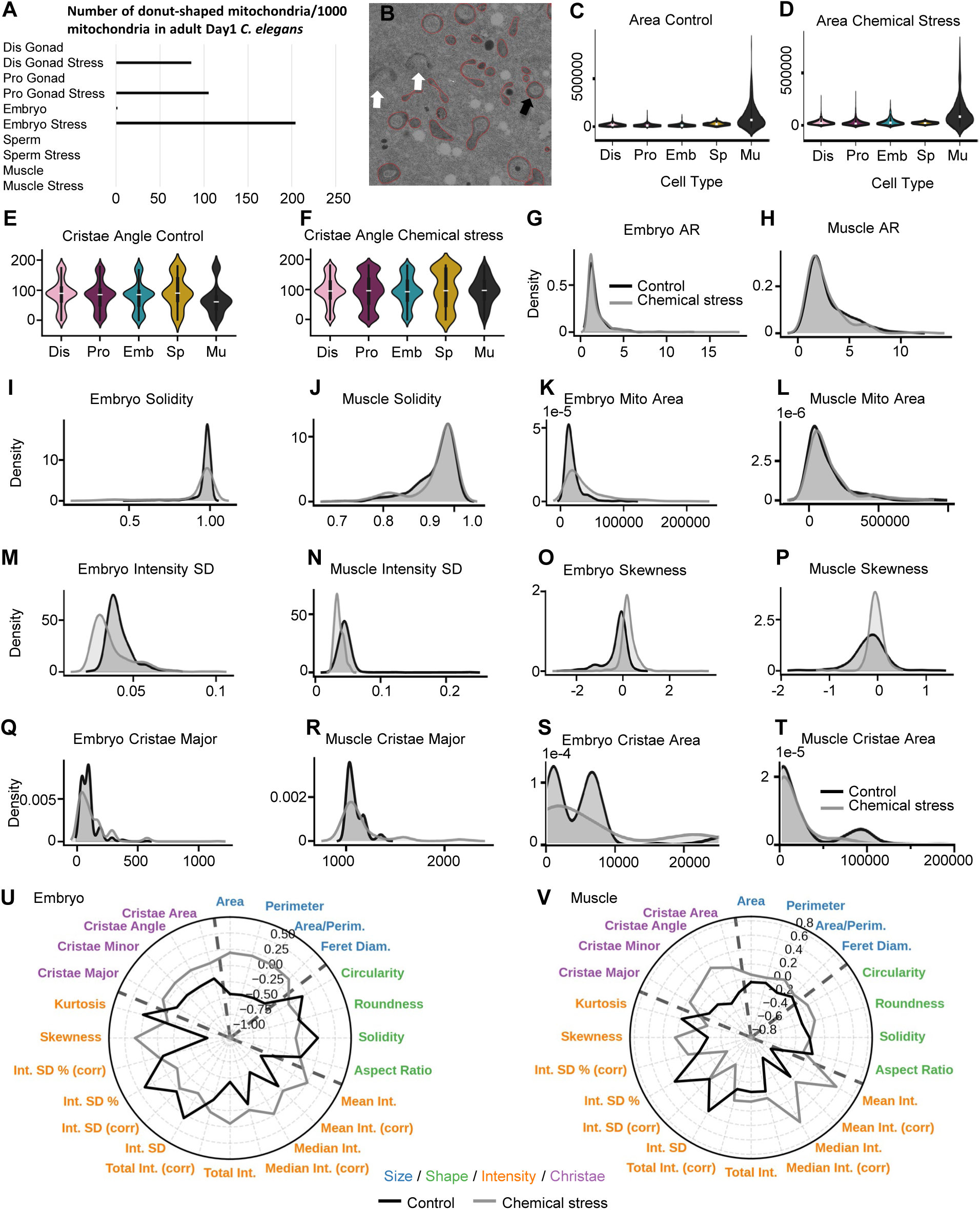
A variety of atypical and donut-shaped mitochondria are formed upon short chemical stress exposure specifically in gonadal and embryonic cells, but not in sperm or muscle cells. A. The number of donut-shaped mitochondria observed in worms exposed (Stress) or not to the mild and short NaN_3_ chemical stress are represented as per thousand mitochondria (‰) in distal gonadal regions, as well as in proximal gonadal regions filled up with oocytes in day1-old-adult worms, in embryos found in the uterus, as well as in sperm cells found in the spermatheca regions or around, and in muscle cells. Total mitochondria number per conditions respectively from top to bottom are: 733; 1998; 745; 1899; 558; 1240; 122; 260; 190; 176. Donut shaped mitochondria, at the most represents 20% (or 200‰) of the total mitochondria in young embryos B. High magnification EM pictures of an oocyte in the proximal gonadal part. The image is overlaid with the automatically segmented and recognized mitochondria, delimited by a red line. White arrows point to the few bended mitochondria that have not been automatically segmented. The black arrow points to donut-shaped mitochondria automatically segmented. Magnification 3000x. C. Mitochondria Area, a metric related to the size of the mitochondria is displayed as violin plots, for all the cell types (Dis: undifferentiated germ cells in the distal gonad; Loop: germ cells in the loop region of the gonad; Pro: oocytes in the proximal gonadal part; Emb: embryonic cells form early embryos in the uterus; Sp: sperm cells; Mu: body wall muscle cells) in control conditions. D. Mitochondria Area, a metric related to the size of the mitochondria is displayed as violin plots, for all the cell types in stress conditions. E. Mitochondria Cristae Angle, a metric related to the cristae orientation of the mitochondria is displayed as violin plots, for all the cell types in control conditions. F. Mitochondria Cristae Angle, a metric related to the cristae orientation of the mitochondria is displayed as violin plots, for all the cell types in stress conditions. G. The Aspect ratio (AR) metric related to the shape of the mitochondria is displayed as density plots, in embryonic cells where there are the most atypical mitochondria after stress exposure (grey line) and to compare it to the control condition (black line). In all density plots from (G-T), the x-axis represents the measurement in its unit (See Table S1 for all metrics units), and the y-axis (Density) represents the proportion of mitochondria per unit of x, with the total area under each curve equaling 100% of the group’s mitochondria. H. The Aspect ratio (AR) metric related to the shape of the mitochondria is displayed as density plots, in muscle cells where there is no appearance of atypical mitochondria after stress exposure (grey line) and to compare it to the control condition (black line) and to the embryonic cells where atypical mitochondria are observed in 20% of the cases I. The Solidity metric related to the shape of the mitochondria is displayed as density plots, in embryonic cells. J. The Solidity metric related to the shape of the mitochondria is displayed as density plots, in muscle cells. K. The Area metric related to the shape of the mitochondria is displayed as density plots, in embryonic cells. L. The Area metric related to the shape of the mitochondria is displayed as density plots, in muscle cells. M. Density plots of the Intensity SD metric related to the Intensity/Texture/Cristae density in embryonic cells. N. The Intensity SD metric related to the Intensity/Texture/Cristae density is displayed as density plots, in muscle cells. O. The Skewness metric related to the Intensity/Texture/Cristae density is displayed as density plots, in embryonic cells. P. The Skewness metric related to the Intensity/Texture/Cristae density is displayed as density plots, in muscle cells. Q. The Cristae Major metric related to the Cristae orientation is displayed as density plots, in embryonic cells. R. The Cristae Major related to the Cristae orientation of the mitochondria is displayed as density plots, in muscle cells. S. The Cristae Area related to the Cristae orientation of the mitochondria is displayed as density plots, in embryonic cells. A zoom has been applied to better visualize the plots, thus the total area under each curve equaling in this case, 90% and 85% for the control and stress group’s mitochondria. T. The Cristae Area related to the Cristae orientation of the mitochondria is displayed as density plots, in muscle cells. A zoom has been applied to better visualize the plots, thus the total area under each curve equaling in this case, 95% and 84% for the control and stress group’s mitochondria. U. Radar plot displaying all 24 mitometrics for both the embryonic cells in control (black) and stress (grey) conditions. Each spoke shows cell-type means, standardized to the common mean (0). The plot center represents the minimum plotted value minus a margin. The metrics can enter four categories, which are color-coded. Metrics categories: size (blue), the shape (green), the Intensity / Texture/ Cristae density (orange), and the Cristae Orientation (purple). V. Radar plot displaying all 24 mitometrics for both the muscle cells in control (black) and stress (grey) conditions. See Table S1 for metrics description and units.

### NaN_3_-induced stress causes global ultrastructural remodeling of cristae across all tissues

Under chemical stress, we applied the EMito-Metrix pipeline to quantify mitochondria across the distal gonad (1,998), proximal gonad (1,899), early embryos (1,240), sperm (260), and body wall muscle cells (176) (**Figure S6**). To accurately segment stress-induced atypical morphotypes (*e.g.,* donut- and horseshoe-shapes), the pipeline underwent an additional fine-tuning cycle. Automated segmentation successfully captured the vast majority of these complex structures, and rare incomplete detections (**Figure 6B**) were manually corrected to ensure accuracy. Because of our extensive sampling scale, statistical power remains robust; any residual undetected mitochondria would merely result in a conservative underestimation of the observed structural differences.

We first performed a global comparison of all 24 mitochondrial morphometrics between control and chemical stress conditions across all cell types (**Figure S3 and Figure S6**). Broadly, baseline tissue-specific distinctions are preserved under stress. For example, somatic muscle mitochondria remain significantly larger than those in all other tissues (**Figure 6C-D**), and the relative differences in most other parameters between cell types remain consistent with control conditions (**Figure S6**). However, global cristae architecture is markedly disrupted. The distributions of all four cristae parameters, most notably the Cristae Orientation Angle, shift significantly following mild sodium azide exposure (**Figure 6E-F**). Specifically, stressed cells exhibit a flattened distribution with a distinct loss of the central peak measures observed in controls, reflecting a widespread reorganization of internal cristae alignment. To dissect tissue-specific responses, we performed pairwise comparisons of individual cell types, such as embryonic cells (which develop donuts) and muscle cells (which do not), under control and stress conditions. While some metrics, such as aspect ratio, remain unchanged across conditions (**Figure 6G-H**), others are significantly altered uniquely in tissues that form atypical mitochondria (distal, proximal gonads, and embryos as shown in **Figure 6I**, **6K**). Specifically, solidity (**Figure 6I-J**), area (**Figure 6K-L**), and to a lesser extent perimeter and circularity (**Figure S6**), shift exclusively in these populations. These quantitative shifts in size and shape metrics perfectly corroborate our qualitative observations of extensive structural remodeling in germ and embryonic cells (**Figure 4 and Figure 5**).

In contrast to these cell-type-specific morphological shifts, chemical stress NaN3 exposure induces global alterations. A notable hallmark of the chemical stress response in almost all measured metrics, beyond the shift in the overall Gaussian profiles, is the presence of consistently longer tails compared to control conditions (**Figure 6G-T**). This pronounced tailing reflects an extensive, wide-ranging heterogeneity within the mitochondrial populations and underscores a systemic loss of regulation. In addition, chemical stress NaN3 exposure induces global alterations across *all* examined cell types specifically in internal architecture. Three distinct classes of parameters are universally impacted: intensity standard deviation (SD) (**Figure 6M-N**), skewness (**Figure 6O-P**), and cristae parameters (**Figure 6Q-V**). These universal shifts reflect widespread cristae disruption and decreased detectable cristae density (indicated by lower intensity SD; **Figure 6M-N**). Notably, somatic muscle mitochondria exhibit equal or greater severity in these internal architectural defects compared to germ and embryonic cells (**Figure 6M-V**). Sperm mitochondria follow this global trend of ultrastructural disruption, though to a lesser degree (**Figure S6**).

Furthermore, stressed mitochondria display broadened peaks in the Cristae Orientation Angle and a loss of multiple spatial frequency peaks in the Cristae Orientation Major (**Figure 6Q-T**). Because a rich conformational repertoire typically produces multiple peaks in healthy mitochondria, this loss signifies a pathological simplification and severe cristae disorganization.

Finally, we leveraged image skewness, a measure of pixel brightness asymmetry relative to the mean, to quantify internal ultrastructural damage. Healthy, well-contrasted cristae typically yield a negative skew (bright pixels with small dark regions), while positive skew reflects dark matrices punctuated by bright highlights (*e.g.,* swollen cristae or large vacuoles). Following stress, skewness increases across all tissues, capturing ubiquitous baseline damage. Remarkably, treated germ and embryonic cells exhibit significantly higher skewness than treated muscle cells (**Figure 6O-P**). Importantly, single-mitochondrion dataset analysis confirms that highly elevated skewness values (≥0.4) specifically correlate with the severe local morphological disruption characteristic of donut-shaped mitochondria and their large, bright internal vacuoles.

### UMAP clustering reveals distinct populations of atypical mitochondria unique to the germline and embryos

UMAP dimensionality reduction further highlights the distinct mitochondrial profiles between control and NaN3-exposed conditions across all examined tissues (**Figure S7**). Strikingly, highly isolated mitochondrial clusters emerge exclusively in the UMAP projections of stressed germ and embryonic cells. These isolated clusters likely represent the distinct atypical morphotypes described above (**Figure 4 and Figure 5**).

Because control and chemically stressed mitochondria segregated so clearly, we applied a multilayer perceptron (MLP) algorithm to automatically predict treatment conditions, based on the calculated mitometrics. Following a training phase, the MLP successfully distinguished stressed from control mitochondria in all cell types. The predictive recall scores/ success rates for control versus stressed conditions were, respectively: 79% and 94% in embryos (**Figure 7A**); 83% and 94% in the distal gonad; 88% and 96% in proximal oocytes; 84% and 91% in sperm; and 98% and 89% in muscle cells.

**Figure 7:**
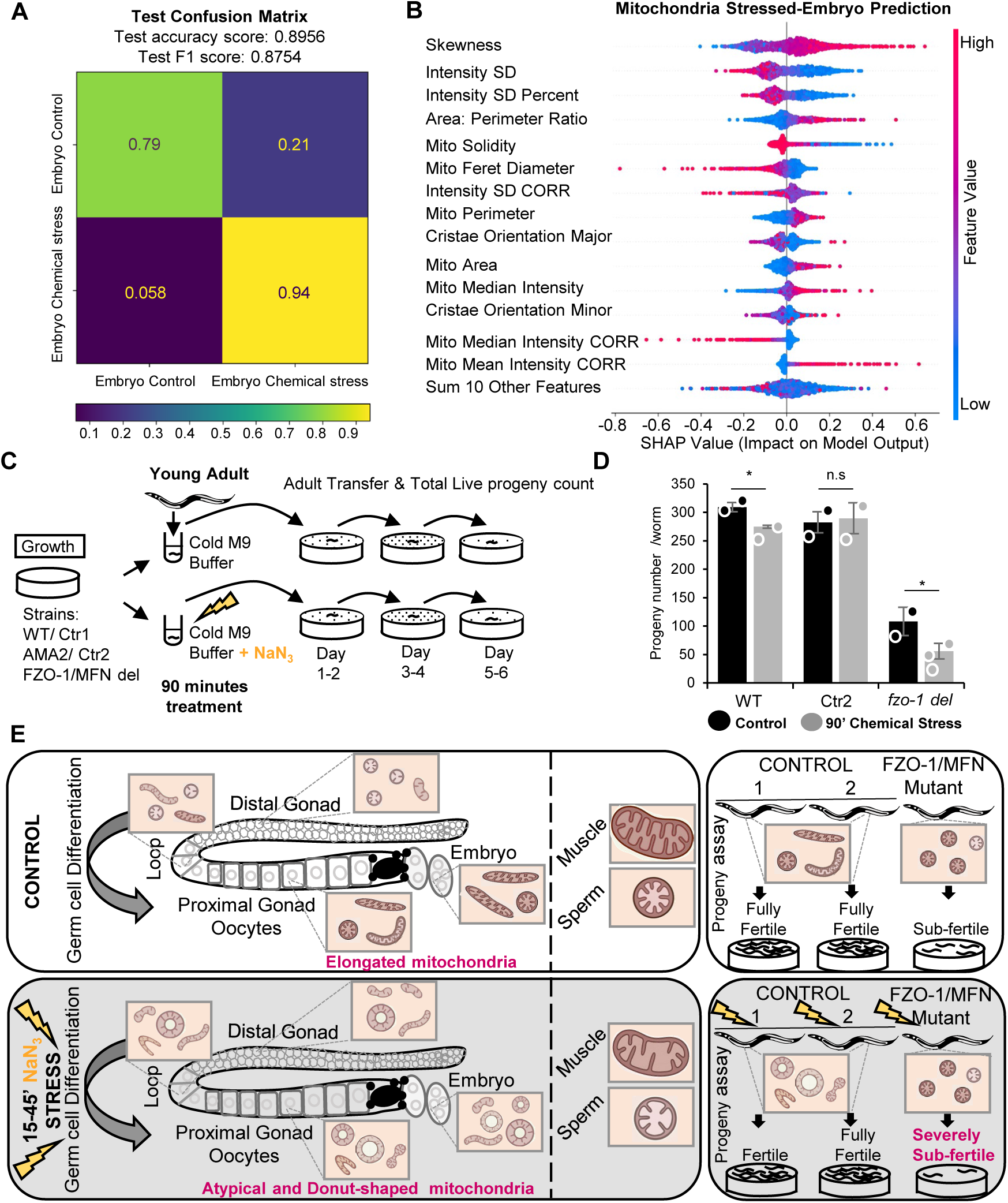
High skewness values reflects donut-shaped mitochondria and fusion-dependent mitochondrial remodeling and donut-shaped mitochondria actively protect female fertility under acute stress. A. Confusion matrix generated by MLP (Multilayer Perceptron) algorithm to predict mitochondria from early embryonic cells in control and stress conditions. B. Global explicability (using MLP algorithm) of the prediction model of the embryonic cells in stress condition compared to the control condition ranking the top-14 most important variables (displayed in order of importance based on the mean of absolute SHAP values). A positive or negative SHAP value on the x-axis means that the variable contributed to the positive or negative estimation of the group prediction. Mitochondria after 15-45 minutes of NaN_3_ exposure exhibit positive skewness within a range and the highest skewness values represent donut mitochondria. C. Schematic view of the progeny/ brood-size assay performed in control conditions and after NaN_3_ exposure. D. Total live progeny per worm (cumulative brood size) for wild-type (Ctr1/WT), control strain (Ctr2/AMA2), and *fzo-1(cjn20)* deletion mutants under control conditions (black bars/circles) and following 90 minutes of acute 30 mM NaN_3_ chemical stress (grey bars/circles). Data represent mean, s.e.m from N = 2-3 independent experiments (individual data points/circles correspond to independent biological replicates). Total number of analyzed worms (n) per condition: WT (n = 14 control, n = 9 stress), AMA2 (n =18 control, n =7 stress), and fzo-1 del (n =15 control, n = 27 stress). p < 0.5, n.s., not significant (Student’s t-test). E. Schematic model of FZO-1-dependent mitochondrial remodeling and fertility protection. Under baseline conditions (top), mitochondria mature from small round profiles into elongated networks with cristae reorganization during germ cell differentiation ensuring normal fertility. Mitochondria from somatic and sperm compartments are distinct. Following acute stress (bottom), germ cells and early embryos specifically form atypical, donut-shaped and spheroid mitochondria. This regulated FZO-1-dependent structural transition acts as a protective defense mechanism vital for preserving female reproductive capacity under metabolic challenge.

We next evaluated the specific morphometrics driving these algorithmic predictions. In embryonic-stressed cells, the highest-contributing predictive features included both the internal ultrastructural metrics altered in all cell-types by NaN_3_ (*e.g.,* skewness, intensity SD) and the macroscopic shape metrics specifically associated with atypical morphotypes (*e.g.,* solidity, area) (**Figure 7B**). Mitochondria after 15-45 minutes of NaN3 exposure exhibit positive skewness within a range and most extreme skewness values represents atypical and donut mitochondria. The strong predictive weight of high skewness further underscores the heavy accumulation of vacuolated, donut-shaped mitochondria in chemically exposed embryos.

Altogether, our findings demonstrate that while NaN_3_-stress exposure induces global alterations in mitochondrial ultrastructure across all observed cell types, the gross remodeling into atypical morphotypes, such as donuts and spheroids, strictly occurs within the germline (pachytene germ cells and oocytes) and early embryos, and is absent from adjacent somatic and sperm cells. This indicates that stress-induced mitochondrial dynamics are governed by a highly specialized, gonad-specific regulatory program.

### Atypical, donut-shaped mitochondria actively protect female fertility under oxidative stress

Atypical, donut-shaped mitochondria have been recently characterized in aging *C. elegans* oocytes, where they depend on the fusion protein FZO-1 to form^19^, as well as in aged mammalian oocytes^12,14–19^. However, because we found these striking morphological changes are rapidly induced and strictly restricted to the gonads and early embryos following acute chemical stress (**Figure 4**, **Figure 7, and Figure S5**), it raises a critical question: do these stress-induced structures only represent collateral damage from oxidative stress, or do they serve as a controlled, active mechanism to protect female fertility?

To determine if these fusion-dependent dynamics play a protective role, we disrupted the mitochondrial fusion. FZO-1, the *C. elegans* ortholog of mammalian Mitofusins (MFN1/MFN2), is an outer membrane GTPase essential for normal fertility, germline mitochondrial elongation^4,13^, and the formation of donut-shaped mitochondria^19^. We reasoned that if these atypical morphotypes actively protect oocytes and embryos during acute stress, animals lacking FZO-1 should exhibit exacerbated fertility defects following chemical exposure.

We exposed young adult worms (at the onset of oogenesis) to 30 mM NaN_3_ for 90 minutes and quantified the total number of live self-progeny (**Figure 7C**). Under standard control conditions, *fzo-1* deletion mutants exhibited the expected baseline fertility defects, yielding an average of 108 progeny per worm, compared to 309 in wild-type (WT) and 282 in a secondary control strain.

Following NaN_3_ stress and a temporary anesthesia-induced delay in egg-laying across all strains^28^, WT worms experienced only a minor 11% reduction in total progeny, while the secondary control strain showed no significant decrease. Strikingly, *fzo-1* mutants suffered a severe 48% drop in fertility, producing an average of just 56 progeny per worm after stress exposure (**Figure 7D**).

Collectively, our results establish that fusion-dependent mitochondrial remodeling in the germline, and the resulting donut-shaped mitochondria are not simply a byproduct of oxidative stress, but an active mechanism shielding *C. elegans* fertility. This raises the compelling possibility that the formation of donut-shaped, spheroid or vacuolated mitochondria represents a conserved evolutionary strategy to protect female gametes from environmental stressors across species. Consequently, the appearance of comparable atypical mitochondria observed in somatic tissues, such as in neurodegenerative diseases, worth investigation for similar protective functions.

## DISCUSSION

In this study, we sought to address two fundamental biological questions: how mitochondrial architecture structurally remodels at the ultrastructural level during continuous gametogenesis, and whether differentiating germ cells respond to acute environmental stress differently than terminally differentiated somatic tissues.

Altogether, our high-resolution transmission electron microscopy (HPF-TEM) and automated EMito-Metrix quantitative morphometric analysis reveal two distinct, highly regulated phenomena. First, under baseline physiological conditions, mitochondria undergo a progressive structural maturation as germ cells differentiate into mature oocytes, transitioning from small, round organelles into elongated, tubular networks featuring more organized and aligned cristae. Second, under acute chemical oxidative stress, these germ cells and early embryos execute a remarkable tissue-specific morphological shift to form atypical donut-shaped and spheroid organelles, whereas adjacent somatic cells and mature sperm remain structurally unchanged. Rather than representing a passive symptom of cellular collapse or organelle damage, our results suggest that this FZO-1-dependent structural reorganization acts as an active, controlled defense mechanism that protects female gametes and preserves reproductive capacity under acute metabolic perturbation (**Figure 7E**).

### Ultrastructural Maturation Prepares the Oocyte for Embryogenesis

Mitochondria of body wall muscles, neurons and several other major cell types have been analyzed via TEM in *C. elegans* in the last decades to different extent^44,45^, however mitochondrial changes during oogenesis have been only recently characterized with fluorescence microscopy resolution^13^. Our TEM quantitative ultrastructural analysis consolidates and found that mitochondrial architecture is progressively remodeled during *C. elegans* gametogenesis, moving from the mitotic tip to the mature proximal gonad, mitochondria shift from small, round forms to elongated, tubular networks. Importantly, it reveals that this longitudinal elongation is tightly coupled with internal cristae reorganization, shifting from a diffuse, directionally disordered cristae orientation in undifferentiated germ cells to a highly aligned, parallel cristae architecture in mature oocytes. This is consistent with a profound metabolic reprogramming and higher ATP production in oocytes^13^, as well as with findings supporting glycolytic metabolism in immature/stem cells vs. mitochondrial OXPHOS in committed/differentiated cells^46^. The structural mitochondrial maturation in oogenesis in *C. elegans* is itself regulated by specific signaling cascades like the MAPK/ERK and sperm-derived MSP pathways^13^, coming from the gonad itself and the surrounding sperm or somatic cells that induce more generally germ cell differentiation and maturation^47–49^.

Because *C. elegans* oocytes are not dormant as in mammals^49,50^ -potentially for decades- and fertilization can occur within minutes to few hours according to their position in the gonad, we propose that these elongated, metabolically primed architectures carry over directly into the early zygotic stages to immediately support the bioenergetic demands of the fertilization, eggshell formation and embryogenesis. Concerning somatic cells, they are already terminally differentiated in our analyzed Day 1-adult samples, highlighting the unique, continuous dynamics of the germline. Mapping the timeline of other cell-types mitochondrial dynamics during differentiation and the exact mitochondrial architecture in a comparative manner of all other cell-types in *C. elegans* remains a key future objective to complete this developmental map. For instance, it would be interesting to determine the sperm mitochondrial rounding and growth during the L4 stage.

### Atypical and Donut-shaped Mitochondria as an Active Defense Mechanism

The emergence of highly complex mitochondrial morphotypes, including donut shapes and spheroids, has historically been often interpreted as a passive hallmark of cellular aging, oxidative damage, or structural degeneration or linked to unknown function^5,14,15,17,18,27,51–53^. However, few recent studies do not fit this traditional view and have implicated donut-shaped mitochondria in osteogenesis, working memory and offspring size control^6,19,54^. Our quantitative ultrastructural and genetic findings fundamentally challenge the passive collapse paradigm, and provide an additional active function for atypical mitochondria, specifically, to defend female germline and early embryos. By demonstrating that fusion-deficient *fzo-1* deletion mutants, suffer a severe 48% decline in total live progeny following acute NaN_3_ exposure, compared to a minor 11% reduction in wild-type animals and no significant decrease in secondary controls, our results suggest that the formation of donut-shaped and spheroid mitochondria is an active, fusion-dependent survival strategy essential for maintaining oocyte quality and fertility.

Conceptually, we propose that these FZO-1-dependent donut shapes^19^ represent an “organelle-level dauer/quiescent state”, a transient, structurally distinct holding pattern that allows mitochondria to remain less active/quiescent and protected during acute environmental perturbation. Much like the organismal dauer diapause in *C. elegans*, which is rapidly deployed under harsh environmental conditions and fully reversed upon the return of favorable cues, this organellar remodeling would function as a temporary biological shield. Importantly, during the worm dauer stage, at the organism-level, the mitochondria even if only partially characterized, are yet different than what we described here in the germ cells under acute stress, with a condensed architecture and a switch to an anaerobic metabolic pathway^44,45,55^.

At the biophysical level, acute exposure to NaN_3_ inhibits Complex IV of the electron transport chain, causing rapid ATP depletion and altered cellular redox balance^28^. We hypothesize that folding into vacuolated spheroid could serve to isolate vulnerable mitochondrial and/or cytoplasmic components, minimize reactive oxygen species (ROS) leakage, sequester mitochondrial DNA (mtDNA), RNAs, proteins, lipids, Ca^2+^ or temporarily shield the organelle from targeted autophagic degradation (mitophagy) until homeostasis is restored^10,11,24,56–59^. By preserving a functional reserve of mitochondrial machinery during metabolic crisis, the oocyte would ensure rapid recovery of oxidative phosphorylation once the stressor ceases.

Future investigations will be necessary to better understand how the protective function take places and to characterize the content of the lumen of the spheroid / donut-shaped mitochondria in *C. elegans* and of the mitochondrial vacuole, and the lumen of the human Mitochondria-Vesicle (M-V) in dormant Human oocytes, to assess the potential similitude of their content as well as the function of those structures. One of many interesting follow-up questions is: Are RNAs found in the lumen, similarly to the function in P granules in *C. elegans* or to the recently identified MARDO, a mitochondria-associated membrane-less compartment that preserves maternal mRNAs^56^ at the M-Ser structures in human oocytes and coordinates their expression during mammalian oocyte maturation? In addition, it is tempting to speculate that under acute, transient environmental stress, the donut conformation could function as a protective biophysical shield, however, if external stress would persist chronically and exceeds the organelle’s buffering capacity, this protective shield would eventually fail, causing these atypical morphotypes to regress and serve only as markers of organelle dysfunction. This kinetic balance is supported by observations in murine embryonic fibroblasts (MEFs), where mitochondrial spheroids form rapidly within 1 hour of CCCP-induced uncoupling, peak between 6 and 16 hours, and completely reverse within hours following stressor removal^27^. The dynamics of those same atypical shapes observed after NaN_3_ exposure and their reversal would be interesting to assess in *C. elegans in vivo*.

### Evolutionary Logic of Stress-Induced Tissue-Specific Remodeling

Remarkably, this remodeling remains confined to specific tissues. Despite systemic exposure of the organism to the chemical agent, massive structural reorganization was observed exclusively within undifferentiated germ cells, maturing oocytes, and early in utero embryos. Adjacent somatic tissues, such as body wall muscle cells, as well as mature spermatozoa harbored within the spermatheca, maintained their baseline mitochondrial architectures without forming atypical morphotypes.

This stark dichotomy reflects the distinct evolutionary imperatives governing the germline versus the soma. The female germline carries the “immortal” genetic and cytoplasmic lineage destined for the next generation, necessitating rigorous organelle quality control and specialized stress-resilience pathways to ensure embryonic viability^24,25,48,57,60–62^. Somatic muscular cells, which prioritize immediate mechanical or bioenergetic output, may lack some of the specialized signaling cascades, yet probably not fully characterized required to execute this rapid membrane bending and donut shaped formation. Furthermore, the inability of mature sperm to undergo this remodeling aligns with their extreme terminal differentiation^37,39,41^. Lacking a canonical nuclear envelope, endoplasmic reticulum, and translationally active ribosomes, mature *C. elegans* spermatozoa are structurally streamlined cells that lack the metabolic and translational machinery required to mount complex organelle-level stress adaptations.

### 2D vs. 3D Interpretations

Obviously, 3D TEM would provide even greater detailed analysis^44^ than 2D TEM images but is more costly in terms of time, money and analysis complexity. Hence, interpreting complex mitochondrial morphotypes requires a rigorous evaluation of 2D TEM limitations. In single 2D sections, a standard tubular mitochondrion cut perpendicularly can appear identical to a small spherical mitochondrion. Consequently, part of the quantitative elongation recorded during germline differentiation in control conditions could theoretically reflect spatial reorientation of elongated mitochondria along the sectioning plane. However, because our quantitative trends directly match the confocal fluorescence-based observations^13^, we are confident that some of the mitochondrial population undergoes true physical elongation.

For the stress-induced donut profiles, analyzing consecutive 80 nm serial sections allowed us to unambiguously confirm that a substantial proportion of these structures represent true 3D mitochondrial spheroids enclose central cytoplasmic lumens. Our serial observations suggest a multi-step morphogenetic pathway: elongated and dumbbell mitochondria undergo localized bending, to form intermediate cup or horseshoe shapes then fuse at initial contact points, and subsequently undergo secondary fusion events to completely or almost completely engulf a cytoplasmic pocket containing ribosomes, cytosol, and occasionally small membranous vesicles. Conversely, serial sectioning consolidated that sperm mitochondria seem strictly spherical in 3D across all planes, as we can interpret from 2D images. Methodologically, our high-pressure freezing and freeze-substitution protocol though provided an advantage compare to more classical TEM studies, with near-native preservation of fine ultrastructural details avoiding the severe organic solvent extraction artifacts that frequently dissolve non-membranous structures like ribosomes, or that frequently prevent to observe adequately atypical mitochondria in conventional chemical fixation protocols^63^. Finally, the EMito-Metrix pipeline provided comprehensive quantitative analysis of mitochondrial ultrastructure, facilitating 2D TEM interpretation and demonstrating broad applicability across species.

### Translational Implications including in Infertility, Reproductive Aging, and Environmental Toxicology

The discovery of protective, donut-shaped mitochondria in *C. elegans* has broad implications for mammalian reproductive biology, stress responses, and aging-related pathologies. Atypical mitochondria, such as vacuolated, donut-shaped, toroidal, and spheroid mitochondria, have now been documented in various organisms across the plant and animal kingdoms^21,22,27,43,64,65^. These include skeletal muscle^20,66^, osteoblasts^6,67^, hepatocytes under long-term alcohol consumption^53^, neurons^52,54^, and oocytes^12,14–19,68^. Interestingly, TEM micrographs in neurodegenerative contexts, such as Parkinson’s disease^51,69^, cognitive aging^54,70^, Huntington’s disease^52^, and Fragile X-associated disorders^71^, explicitly depict donut and spheroid mitochondrial topologies, which frequently correlate with local metabolic stress levels and cellular dysfunction. In light of our functional findings, some of these atypical mitochondrial structures warrant functional re-evaluation. Rather than acting as passive correlates of pathology, they may represent an evolutionarily conserved, adaptive defense mechanism that preserves cellular viability during metabolic stress.

Mitochondrial plasticity generally supports oogenesis in vertebrates and mammals^50^. Interestingly, donut-shaped and other atypical mitochondrial structures are reported in oocytes across various vertebrates and mammals, ranging from Xenopus, hamsters, cows, sheep, goats, and mice to humans^14,16–18,60,68,72^. These structures include cap-shaped or hooded mitochondria, dumbbell, ring, and horseshoe morphotypes, alongside vacuolated mitochondria where cytoplasmic content and ribosomes fill the vacuoles, as interpreted from early electron microscopy tomographs^12,14,16,18^. In most species, these profiles closely resemble the atypical mitochondria observed in aged^19^ or stress-challenged *C. elegans* oocytes (Figures 3–6), and similarly to *C. elegans*, they become more prevalent during later stages of oocyte development or with age^10,68,73^. While mouse and human oocytes display subtle differences in atypical morphologies compared to other mammals and each other^12,14,16,18^, human growing oocytes also display numerous round-to-oval shaped mitochondria containing a large internal clear vacuole, alongside a paucity of arched cristae and poorly developed cristae^12,14,15,68,72^. These features suggest low, quiescent catalytic activity, supported by declining mitochondrial-coupled oxygen consumption during human oocyte development^60,74^. Interestingly, atypical and even these typical human oocyte mitochondria share common features with *C. elegans* donut-shaped mitochondria under oxidative stress conditions. Therefore, we hypothesize that human atypical and vacuolated mitochondria may play a protective role and represent a healthy marker in growing oocytes which might be under oxidative stress, environmental challenges or other specific cues. Conversely, their persistent presence post-ovulation or post-fertilization might indicate deficient energy supply and compromised bioenergetic capacity^15^.

Building on this vulnerability to metabolic stress, oocytes exhibit heightened susceptibility to environmental toxins, including heavy metals, industrial pollutants, pesticides, and consumer chemicals^75,76^. Specifically, exposure to perfluorononanoic acid (PFNA), a widespread industrial antifouling and water repellent agent, or methyl parahydroxybenzoate (MP), a common preservative in cosmetics and foods, significantly elevates reactive oxygen species, impairs MAPK/ERK signaling, disrupts meiotic spindle assembly, and collapses mitochondrial membrane potential, culminating in maturation failure and infertility^75^.

Given the conserved nature of mitochondrial dynamics during oogenesis across species and the quantifiable formation of stress induced donut mitochondria detectable also via live fluorescent imaging^13,19^, we propose the *C. elegans* gonad as a valuable, prospective high throughput *in vivo* model for reproductive aging, environmental toxicology, and pharmacological discovery. Free from ethical restrictions and high maintenance demands of mammalian models, *C. elegans* offers potential accessibility for tracking oocyte lineage and mitochondrial fate *in vivo*. This system could enable precise titration of acute and chronic chemical exposures to screen protective compounds that neutralize oxidative stress, restore mitochondrial membrane potential, and oocyte damage^75^. Furthermore, utilizing *C. elegans* to elucidate the molecular mechanisms governing protective mitochondrial structures may help establish novel criteria to assess oocyte competence and identify donor eggs with superior adaptive resilience for *in vitro* maturation or *in vitro* fertilization applications, addressing age related declines in oocyte quality. Crucially, we envision that this proposed *C. elegans* gonadal mitochondrial model could allow researchers to evaluate toxicity, therapeutic efficacy, and oocyte competence across six integrated biological scales: molecular, organelle, cellular, tissue, whole organism, and intergenerational outcomes, including brood size, egg laying kinetics, and offspring stress resilience. Leveraging this model could advance fundamental understanding of mitochondrial structural plasticity, aid in optimizing maturation media for human *in vitro* fertilization^77^, complement research into Mitochondrial Replacement Therapy^75^, and unlock therapeutic targets to extend reproductive longevity and combat age related fertility decline, a major public health concern that has increased by 84 percent globally since 1990^76^.

## MATERIAL AND METHODS

### Worm growth conditions and Strains

We followed standard procedures for maintaining *C. elegans* strains. Worms were grown on Nematode Growth Medium (NGM) plates seeded with *E. coli* OP50 strain and maintained at 22.5°C in a cooling incubator. When experimented on, worms were manipulated in a temperature-controlled environment with temperature around 22°C (± 2°C). The following nematode strains are available from CGC and were utilized in this study: N2, wild-type Bristol isolate; BXN723, *fzo-1(cjn20*) II^78^ and CA1200 *ieSi57 [eft-3p::TIR1::mRuby::unc-54 3’UTR + Cbr-unc-119(+)] II*^79^. Additionally, strain AMA2 was used as a control. Although AMA2 was originally constructed for a separate project, auxin was omitted from all experiments in this study; consequently, no targeted protein degradation was induced, making it a suitable control. Strain AMA2 has the genotype *ieSi57* [*eft-3p::TIR1::mRuby::unc-54* 3’UTR + *Cbr-unc-119* (+)] II; *lsm-8(amy1*[*lsm-8*::linker::AID::3xFLAG]) IV. The *amy1* allele was generated via CRISPR-Cas9 genome editing as described previously^80^. The strain was subsequently outcrossed four times with N2, crossed with CA1200, and outcrossed twice more. The gRNA used was 5’-aatccaaattggtgcaagt-3’ and the gblock sequence inserted to fuse to LSM-8 C-term was 5’-cgatgaagaactcgacaaacgcgttgatctgg aaaatgtgaaagcggctccactcgctccaatctggattccccaaATGGATGAACCACCATCTAAGAAGTCTCGCGCTGATAGC atgcctaaagatccagccaaacctccggccaaggcacaagttgtgggatggccaccggtgagatcataccggaagaacgtgatggtttcctgcc aaaaatcaagcggtggcccggaggcggcggcgttcgtgaagATGGATTACAAGGATGACGATGACAAGGACTACAAGGATG ATGACGATAAGGATTACAAGGATGATGATGATAAGtaatttccacacaaatcgatattctgcattgacgcgttgtattgactggtt attc-3’.

### TEM Sample Preparation

Transmission electron microscopy (TEM) was performed on synchronized Day 1 (D1) gravid adult animals. Age synchronization was achieved by manually picking young adult (YA) hermaphrodites approximately 18 hours (ON) prior to treatment, identified based on vulval morphology, body length, and the absence of eggs in the uterus. Synchronized D1 gravid adults of the AMA2 strain were harvested and incubated in ice-cold M9 buffer either in the presence (chemical stress condition) or absence (control condition) of 30 mM (0.2%) sodium azide NaN_3_.

### High Pressure Freezing and Transmission Electron Microscopy

For cryofixation, worms were handpicked and placed between 3 mm gold-coated copper carriers within a mix of 20% BSA solution and an OP50 bacterial paste (type A and B, 200 µm deep, Leica Microsystems). The sandwiched samples were frozen using a Leica EM ICE high pressure freezer (Leica Microsystems, Austria). After freezing, the samples were immersed in a substitution medium consisting of 2% osmium tetroxide, 0.5% glutaraldehyde, and 0.1% uranyl acetate in absolute acetone, and pre-cooled to -90 ◦C using an Automated Freeze-substitution System (Leica Microsystems, Germany). The AFS program was run as follows: −90°C for 107 h, −90°C to −60°C for 6 h (5°C/h), −60°C for 8 h, −60°C to −30°C for 6 h (5°C/h), −30°C for 8 h, -30 to 0°C for 6 h (5°C/h), 0 to +15°C for 15min. After freeze-substitution, the samples were washed three times with absolute acetone at room temperature and embedded in EMBed 812 epoxy resin. After 48 h of polymerization at 60 °C, ultrathin sections (80 nm thick) were mounted on single-slot Formvar-coated copper grids. Finally, sections were stained with Uranyless and lead citrate. Grids were examined with a TEM (Jeol JEM-1400, JEOL Inc, Peabody, MA, USA) at 80 kV. Images were acquired using a digital camera (Gatan Rio9, Gatan Inc, Pleasanton, CA, USA).

### Image Analysis

The various stages of the image analysis -image preprocessing, mitochondria segmentation, morphological measurements and data prediction-were carried out using EMito-Metrix software, which segments individual mitochondria and 28 morphological and ultrastructural descriptors per mitochondrion^26^ (https://github.com/Restore-Lab/EMitoMetrix).

#### 1- Batch segmentation & Morphological analysis

The raw TEM images used for the EMito-Metrix analysis were all taken at a magnification of 3000x, in which a pixel represents 2.2 nm. Images with a 1 µm scale bar in Figures 1-2; 4–5 are examples of images at a magnification of 3000x. The raw TEM images were first normalized on amplitude (normalization relative to the mean background intensity of the image) to attenuate global gray level variations observed between images than may be related to variations in the amount of staining agent. Automatic mitochondrial segmentation was then performed using the Cellpose BIOP wrapper^81^ implemented in the EMitoMetrix tool.

The parameters taken as input to the plugin and set by the experimenter are the type of prediction model (Species-specific *C. elegans* model) and the diameter (in pixels) of the objects to be segmented. Species-specific *C. elegans* model has been used here to segment mitochondria from the EM images. The mitochondrial segmentation maps (or labelled maps) generated as output of EMito-Metrix were then used in the same tool to calculate the various individual (per mitochondrion) or average (per image) morphological and ultrastructure measurements. EMito-Metrix outputs were used unmodified, with one tab-separated file per imaged field of view, one row per mitochondrion. The dataset analyzed here comprises 558 files and 8 414 individual mitochondria, covering six cell types (distal gonad, loop, proximal gonad, embryo, sperm, body-wall muscle) under two conditions (control and chemical stress).

#### 2- Data display and prediction

The morphological and ultrastructure measurements from the image analysis were then analyzed using data display and data prediction modules as implemented in the EMitoMetrix tool. Data display was used to generate graphical outputs from mitochondrial measurements, and compare these measurements between the different conditions. Those downstream analysis was performed with mito-marker v0.1.1, a Python package developed for this project, under Python 3.12.1 with anndata 0.12.10, NumPy 2.4.2, pandas 2.3.3, SciPy 1.17.1, scikit-learn 1.8.0, umap-learn 0.5.11, SHAP 0.50.0, Matplotlib 3.10.8 and seaborn 0.13.2. Measurements were loaded into a single AnnData object in which each row is one mitochondrion, the 28 EMito-Metrix descriptors form the data matrix, and cell type, condition, source image and specimen are stored as per-mitochondrion annotations. Four descriptors that encode a mitochondrion’s position within the field of view rather than its morphology (Mito_CentroidX/Y, Mito_FeretX/Y) were excluded from all analyses, leaving 24 analytical descriptors. Descriptors were grouped into four families for display - size, shape, intensity, and cristae orientation- and this grouping is used consistently in every figure in which multiple descriptors appear. The description of the different mitochondrial metrics is provided in the **Supplemental Table 1**.

We used the methods Uniform Manifold Approximation and Projection (UMAP) for dimensionality reduction analysis of these 24 mitochondrial descriptors.

From data prediction module, we used a Neural Networks with Multi-Layer Perceptron (MLP) in order to evaluate the performance of the model in classifying mitochondria as either from the distal or proximal part in control conditions, or mitochondria found in the control or chemical stress conditions. To identify the main features used by the models to perform its prediction, the Shapley Additive exPlanations (SHAP)^82^ post-hoc local explainability method has been used. It uses a game theory framework to compute the contribution of each feature to the prediction of each instance, for the model. The contributions are then aggregated on a single plot to identify the contribution of each feature at a glance. Violin plots and density plots show raw, unnormalized measurements in the units produced by EMito-Metrix. For the radar plots, UMAP embeddings and classification models, each descriptor was standardized independently across the mitochondria included in that comparison (z-score: mean subtracted, divided by the standard deviation), so that descriptors spanning different units and ranges contribute comparably.

Violin plots show the full distribution of a descriptor per condition, with an inner box plot marking the median and interquartile range. Density plots show Gaussian kernel density estimates (SciPy gaussian_kde, Scott’s rule for bandwidth). Radar plots show, for each descriptor, the mean of the standardized values per condition; each spoke is scaled independently to its own 1st-99th percentile range across the mitochondria plotted, so that descriptors of different magnitude remain comparable on one axis.

### Live Progeny Assay

Total reproductive output was determined using a standard *C. elegans* brood size assay adapted for post-treatment recovery. Age-synchronized young adult (YA) hermaphrodites of the wild-type (WT) N2 strain, the AMA2 control strain, and the *fzo-1(cjn20)* mutant strain (BXN723) were selected according to the same criteria as above. Worms were picked and transferred into 30 µL of ice-cold M9 buffer supplemented with or without 30 mM (0.2%) sodium azide NaN_3_ in 60-well conical minitrays (Cat. No. 12-565-67) and incubated for 90 min. Following treatment, the supernatant was gently exchanged three times with fresh M9 buffer using a fine glass Pasteur pipette, ensuring worms remained at the bottom of the well. Animals were subsequently transferred onto individual NGM plates seeded with *E. coli* OP50, and residual liquid was carefully aspirated to eliminate remaining NaN_3._ The initial number of transferred adults was recorded. To determine reproductive capacity, surviving adults were transferred to fresh seeded NGM plates every 24–48 hours throughout their egg-laying period (5–6 days total). Following adult removal, plates were incubated for a sufficient time to allow all laid eggs to hatch, and live larvae (F1) were enumerated promptly prior to reaching reproductive maturity to avoid counting second-generation (F2) progeny. Larval counts were normalized to the number of live adults present during the corresponding lay window. The total live progeny count per hermaphrodite was calculated as the cumulative sum of live larvae produced across the entire reproductive lifespan.

## ACKNOWLEDGMENTS

We appreciate and are grateful to Iskra Katic for her help for injection, to Merav Cohen for her help with the manuscript, and to technical assistance from Sabrina Babouram, Valerie Nicolas, Alixe Mauchien Battistetti for worm culture media. We thank the CMEAB platform for preliminary tests. We thank team members and colleagues for the discussions and feedbacks on the manuscript. We are grateful to the reviewers and editors for their precious time.

This project and A.M research team was supported by the French National Research Agency (JCJC, ANR-21-CE12-0011). The METi imaging facility (Genotoul-TRI), member of the national infrastructure France-BioImaging supported by the French National Research Agency (FBI BIOGEN, ANR-24-INBS-0005. The RESTORE data analysis facility (Genotoul-TRI) member of France-BioImaging is supported by the French National Research Agency (ANR-24-INBS-0005 FBI BIOGEN). J-PP is a member of the IHU HealthAge, supported by the French National Research Agency (ANR) as part of the France 2030 program (ANR-23-IAHU-0011). GS is supported by the Agence Nationale de la Recherche for the grant EUR CARe (ANR-18-EURE-0003). Some strains were provided by the CGC, which is funded by NIH Office of Research Infrastructure Programs (P40 OD010440). We also thank the CGC European Hub.

## AUTHOR CONTRIBUTIONS

A.M conceived the study. V.S, A.M, S.B, A.D and M.G. performed experiments and S.B, V.S and A.M contributed to the experimental design. A.M, carried out image acquisition and image analysis and M.V carried the EMito-Metrix pipeline training and analysis. G.S carried data visualization for some plots. A.M, M.V and JP-P contributed to the interpretation of results. A.M. wrote the first draft of the manuscript, and all authors contributed to the final version.

---

**Figure S1:**
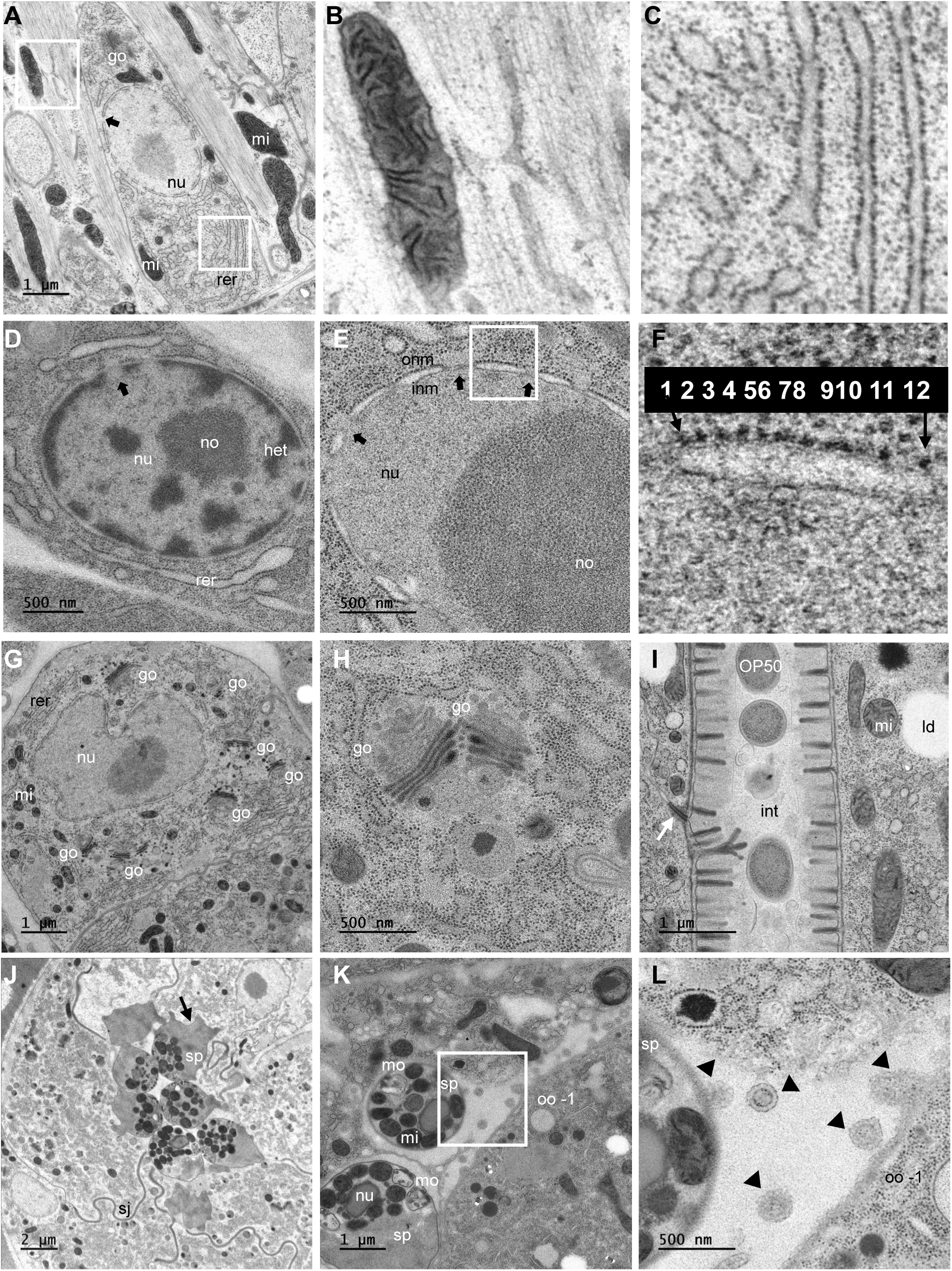
Transmission electron microscopy with cryo-fixation that result in highly resolved images of a Adult Day 1 *C. elegans*. A. Transmission electron micrograph (TEM) of a longitudinal view of a cell in the pharynx region with smooth muscle fibers surrounding it. The series of highly resolved and contrasted pictures we have acquired allow us to observe mitochondria which are the grey-dark stained organelles, one of them in the upper inset is enlarged in (B), where we can clearly observe the ultrastructure including the cristae inside the mitochondria; the Rough Endoplasmic Reticulum (rer) which is largely present in that cell-type and the inset is enlarged in (C) where the ribosomes -black small dots-coat the endoplasmic reticulum. In this cell (a) a golgi apparatus (go) is visible as well as the nucleus (nu) in its approximate mid-section, where the double outer and inner nuclear membranes are visible and interrupted by the nuclear pore complexes (NPCs), one example is shown with the black arrow. B. Mitochondria enlarged from (A). C. Rough ER enlarged from (A). D. High magnification of a nucleus in which the heterochromatin (the darkest matter; het) is localized at the nuclear periphery and in patches in the nucleoplasm, the nucleolus (no) is the roundish part stained a bit lighter than the heterochromatin inside the nucleus. The heterochromatin as expected can localize beneath the nuclear envelope (NE) but excluded from regions where NPCs are localized (black arrow for example). Rough Endoplasmic Reticulum (rer) is observable in close proximity to the NE. E. High magnification of another cell type nucleus in which the outer nuclear membrane (onm) and inner nuclear membrane (inm) fuse at the NPCs (black arrows). no: nucleolus. In the white square a portion of the NE in between two NPCs is enlarged in (f), where the onm is decorated with ribosomes, as expected. F. The resolution of the picture allows to identify the number of ribosome -black small dots-on portions of the onm in between two NPCs. In this example we can count 12 ribosomes. G. Transmission electron micrograph of a different cell type, enriched with golgi apparatus (go). H. High magnification of two golgi organelles one close to each other surrounded by rough endoplasmic reticulum. I. Transmission electron micrograph (TEM) of a longitudinal view of the lumen portion of the intestine of *C. elegans*. Inside the lumen, 3 or 4 E. coli OP50 bacteria can be seen, one of the four in that picture seem to have been lysed whereas the 3 others seem relatively intact with bacterial ribosomes detectable internally. One microvilli layer homogeneous in length is detectable on both side of the intestinal lumen. A junction (white arrow) is detectable outside the apical membrane. Lipid droplets (ld) and mitochondria (mi) are found in the intestinal cytoplasm among other organelles. Bacteria and mitochondria can have similar sizes, but their ultrastructure is clearly different. J. Micrograph of valve cells from spermatheca region with nicely organized pleated septate junctions (sj) and 6 spermatids (sp) with pseudopod visible (black arrow) for some of them. K. Transmission electron micrograph of two sperm cells in proximity to the proximal oocyte. The dark grey organelles are mitochondria and the white structured organelles are membranous organelles (mo). (nu) represents the condensed genome found in spermatids, where the nuclear compartment is hardly definable given the lack of nuclear envelope in this cell-type. Thanks to the well-preserved membranes and structures in our samples, we can notice in the white boxed region enlarged in (l) an exchange of extracellular vesicles (EVs) between the sperm cells (sp) and the proximal oocyte (oo-1), similarly to previously shown vesicles that transport Major Sperm Proteins (MSPs) that contribute to oocyte maturation. L. Zoom in on exchange of extracellular vesicles (EVs) from a spermatocyte (sp) to the proximal oocyte (oo -1) from panel (K).

**Figure S2:**
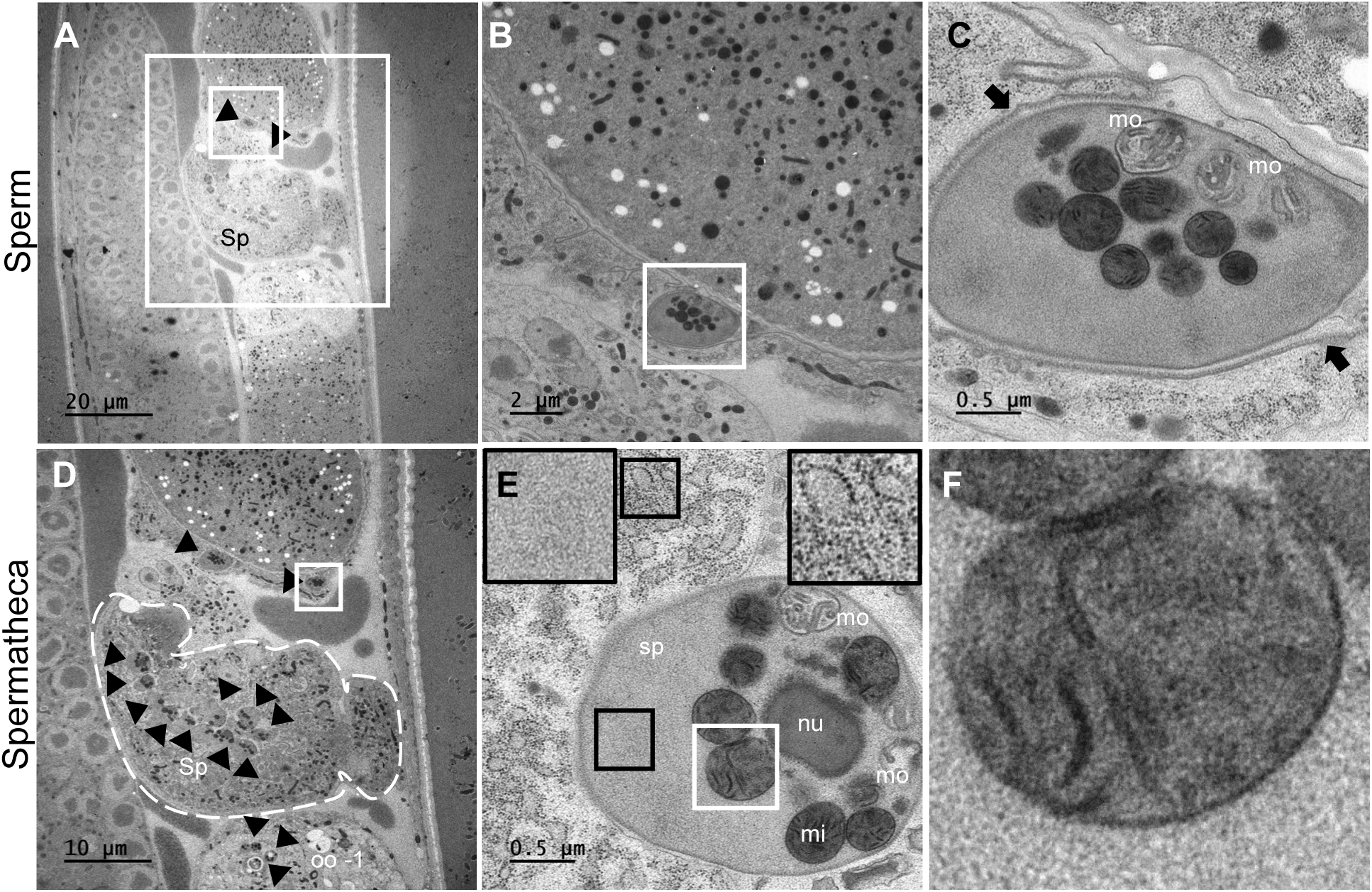
Mitochondrial morphology in spermatids and ultrastructure of *C. elegans*. A. Transmission electron micrograph (TEM) of a longitudinal view of the spermatheca (Sp). The spermatheca is located in between the last oocyte (-1, bottom part) and the first fertilized egg (upper part), and the spermatheca is juxtaposed to the distal gonad (left part) on that image. Two spermatocytes (black arrowheads) are found in proximity to the first egg The region marked by a small square is enlarged in (B), the bigger square is enlarged in (D). B. Zoom in of one of the sperm cell from (A). C. The grey organelles are mitochondria and the white structured organelles are the membranous organelles (mo). Black arrowheads might represent the vitelline layer of the embryo eggshell which envelope this sperm cell. D. The spermatheca (Sp) delimited by a dashed white line. The spermatheca is located in between the last oocyte (oo -1, bottom part) and the first egg (upper part), and the spermatheca is juxtaposed to the distal gonad (left part) on that image. Sperm cells (black arrowheads) are observed in the spermatheca as well as in proximity to the oocyte -1 and to the first(s) embryo(s) in all worms we could image the spermatheca and the oocyte and embryos surrounding it. The region marked by a square is enlarged in (E). E. Enlargement of the region marked by a square in (D). The grey organelles are mitochondria (mi) and the white structured organelles are the membranous organelles (mo). We can notice in the black inserts enlarged in the upper corners that in that sperm cell (sp; as well as in all other sperm cells observed with high magnification and resolution) no ribosomes (back dots) or any endoplasmic reticulum types are detectable in the cytoplasm of the spermatids (left upper insert) in contrast to the cytoplasm of other cell types (right upper insert) surrounding the spermatids. The halo at the periphery of the nucleus (nu) of the spermatid and the absence of nuclear membranes as already characterized are also clearly visible. The region marked by a white square is enlarged in (F). F. Zoom in on one of the mitochondria from (E).

**Figure S3:**
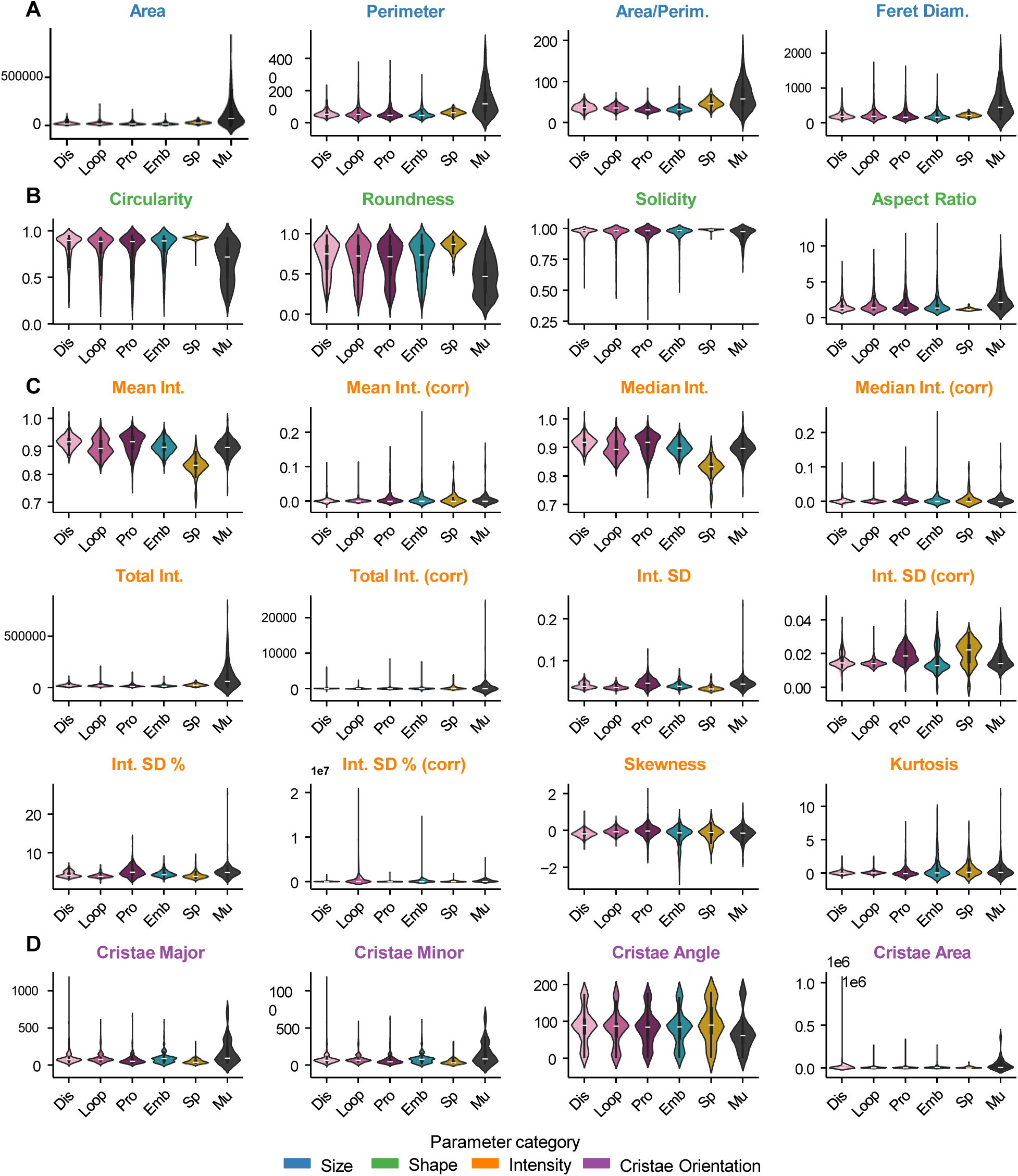
EMito-Metrix pipeline, all 24 metrics displayed as violin plots for all cell types in the control condition. A. Mitochondria metrics related to the size of the mitochondria (Area, Perimeter, the Ratio of the Area/ Perimeter and the Feret Diameter) are displayed using violin plots for mitochondria of all cell types (Dis: undifferentiated germ cells in the distal gonad; Loop: germ cells in the loop region of the gonad; Pro: oocytes in the proximal gonadal par; Emb: Embryomic cells form early embryos in the uterus; Sp: Sperm cells; Mu: body wall muscle cells ) in the control condition. B. Mitochondria metrics related to the shape of the mitochondria (Circularity, Roundness, Solidity and Aspect Ration) are displayed using violin plots, for all cell types. C. Mitochondria metrics related to the Intensity of the mitochondria (Mean Intensity, Mean Intensity Corrected, Median Intensity, Median Intensity Corrected, Total Intensity, Total intensity Corrected, Intensity Standard Deviation, Intensity Standard Deviation Corrected, Intensity Standard Deviation Percent, Intensity Standard Deviation Percent Corrected, Skewness, Kurtosis) are displayed using violin plots, for all cell types. D. Mitochondria metrics related to the Cristae Orientation of the mitochondria (Cristae Orientation Major, Cristae Orientation Minor, Cristae Orientation Angle, Cristae Area) are displayed using violin plots, for all cell types. See Table S1 for metrics description and y-axis units.

**Figure S4:**
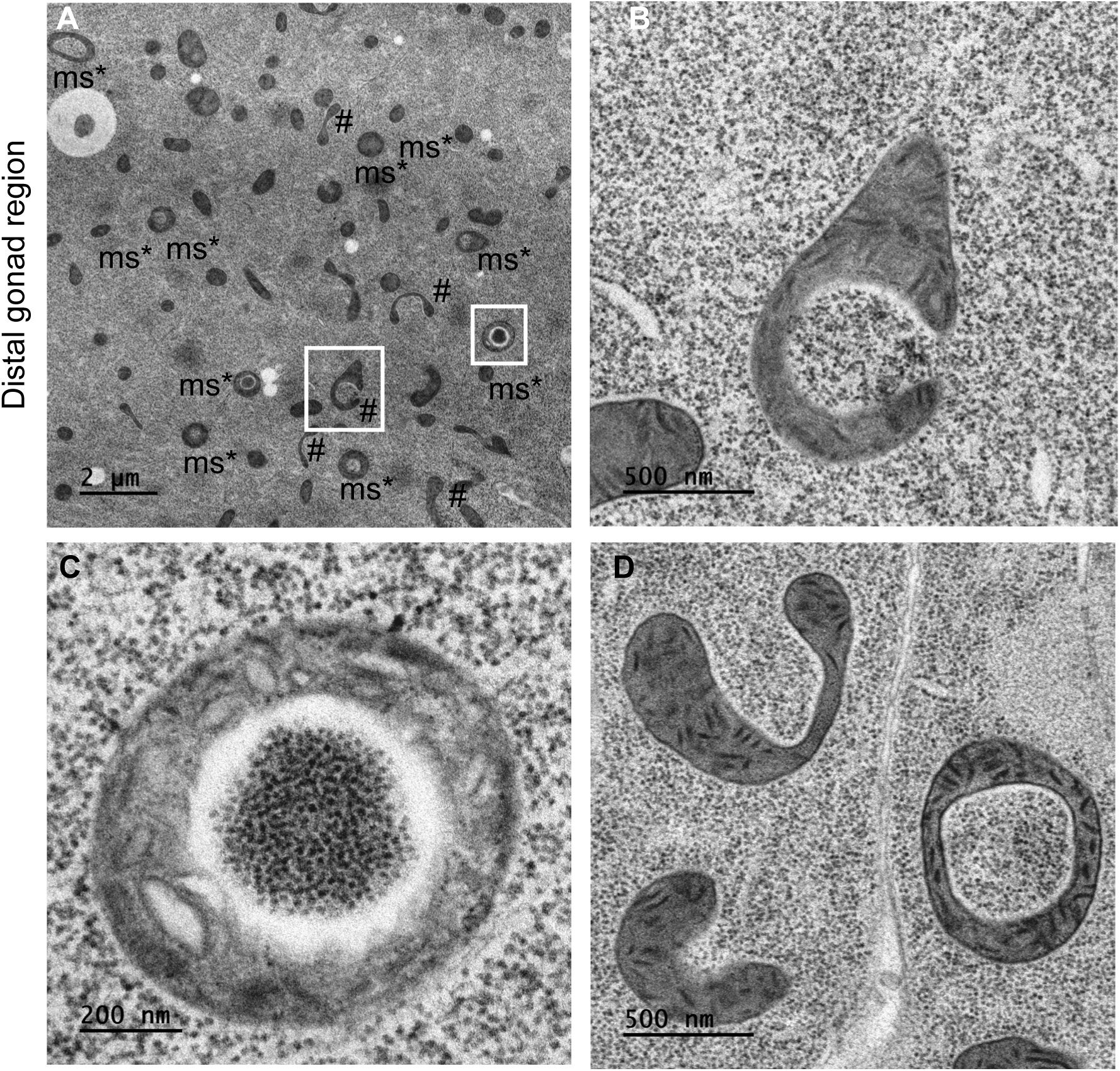
Mitochondrial morphology and ultrastructure in distal gonads in anesthetized *C. elegans* Adult Day 1. A. Transmission electron micrograph (TEM) of a longitudinal view of a part of the distal gonad. The numerous grey round organelles are mitochondria. Many of them have a spheroid shape (ms), or called donut-shaped or ring-shaped mitochondria, other mitochondria have elongated #, horseshoe or dumbbell shapes. B. Zoom in on one of the mitochondria from (A), with example of elongated bended mitochondria, with a hook shape. C. Zoom in on one of the mitochondria from (A), with an example of a donut-shaped mitochondria, which is probably a spheroid mitochondrion with an internal lumen or vacuole in 3D, including probably cytoplasmic ribosomes. D. Zoom in other mitochondria representing elongated and bended mitochondria, like horse shoe-shape mitochondria and a donut shaped one.

**Figure S5:**
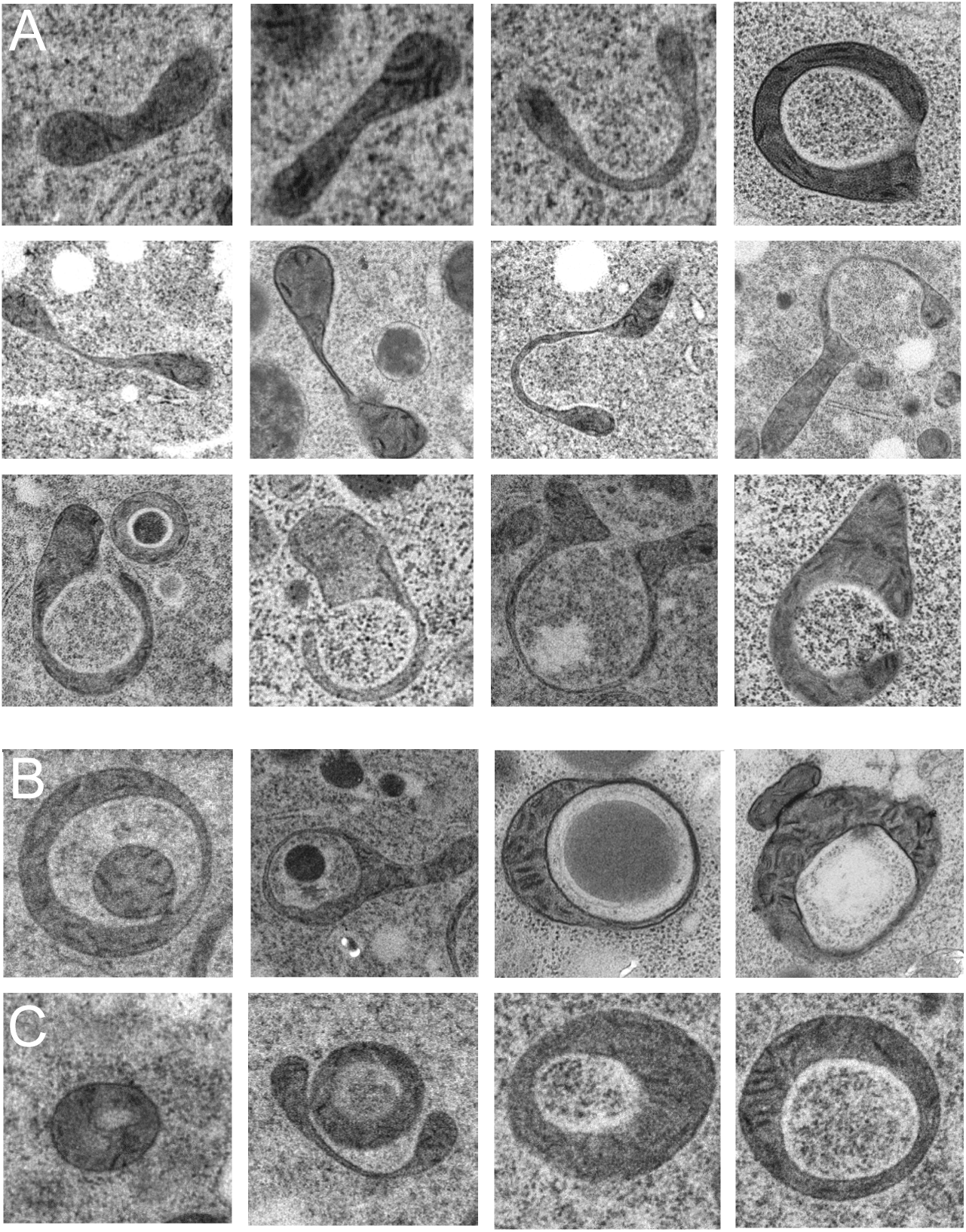
Atypical mitochondrial morphology in anesthetized gonads of *C. elegans*. A. Transmission electron micrographs (TEM) of some examples of elongated shaped mitochondria with dumbbell shape, horseshoe, hook or other shapes. B. Transmission electron micrographs (TEM) of some examples of spheroid / donut shaped mitochondria (ms*). Various ms* mitochondria seem to embed in addition to ribosomes and cytoplasm other cytoplasmic constituents such as portion of mitochondria, of yolk particles, lysosomal or lipid droplets. C. Mitochondria with various ratios between the area sizes of the internal light matter and the area size of the mitochondria (often in the same cells) suggest that at least some of the observed spheroid / donut mitochondria ms* are 3D spheroid mitochondria filled inside with some cytoplasm including cytoplasmic ribosome content.

**Figure S6:**
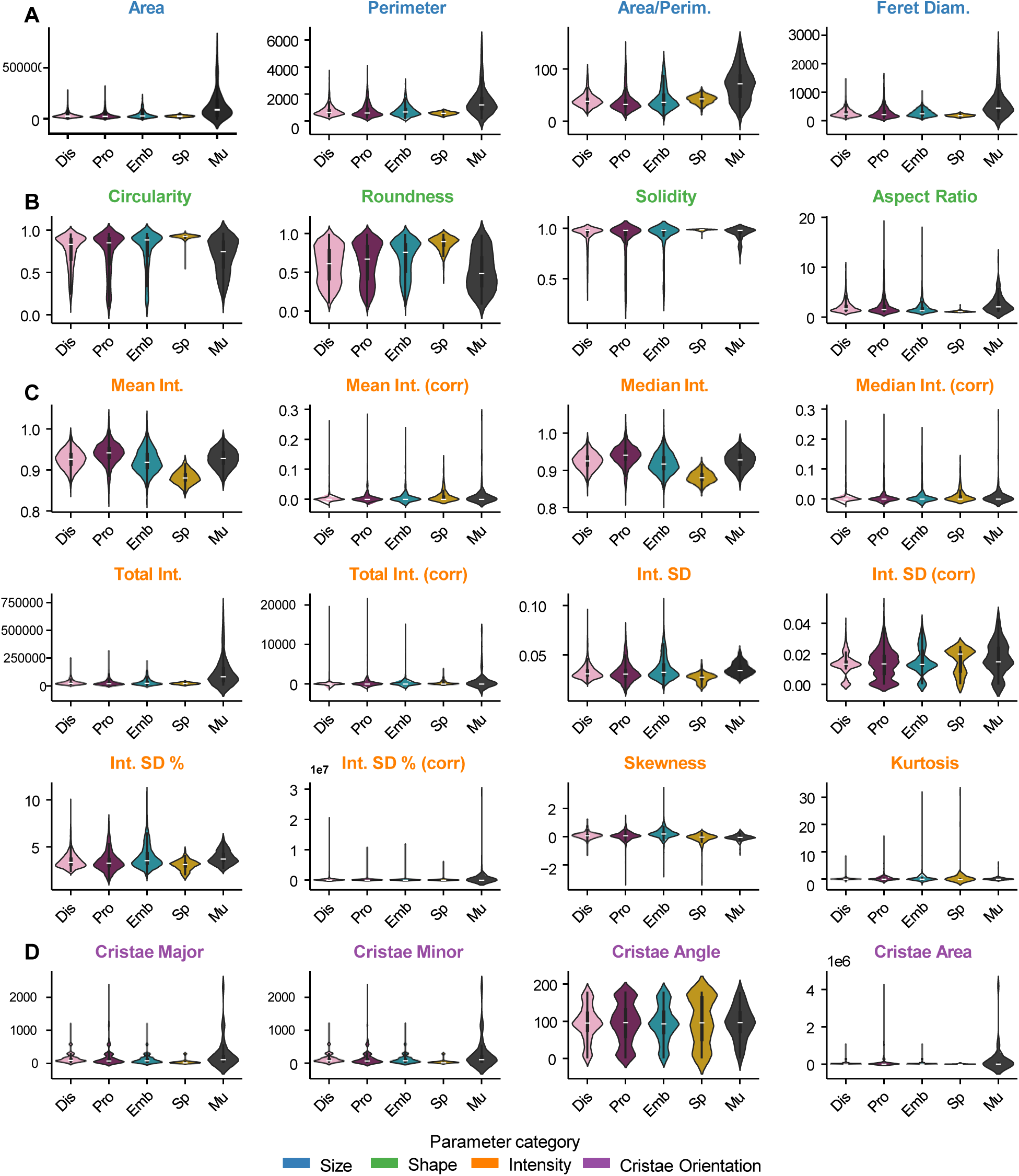
EMito-Metrix pipeline, all 24 metrics displayed as violin plots for all cell types in the “Chemical Stress” condition. A. Mitochondria metrics related to the size of the mitochondria (Area, Perimeter, the Ratio of the Area/ Perimeter and the Feret Diameter) are displayed using violin plots for mitochondria of all cell types (Dis: undifferentiated germ cells in the distal gonad; Loop: germ cells in the loop region of the gonad; Pro: oocytes in the proximal gonadal par; Emb: Embryomic cells form early embryos in the uterus; Sp: Sperm cells; Mu: body wall muscle cells ) after short stress exposure. B. Mitochondria metrics related to the shape of the mitochondria (Circularity, Roundness, Solidity and Aspect Ration) are displayed using violin plots, for all cell types, after short stress exposure. C. Mitochondria metrics related to the Intensity of the mitochondria (Mean Intensity, Mean Intensity Corrected, Median Intensity, Median Intensity Corrected, Total Intensity, Total intensity Corrected, Intensity Standard Deviation, Intensity Standard Deviation Corrected, Intensity Standard Deviation Percent, Intensity Standard Deviation Percent Corrected, Skewness, Kurtosis) are displayed using violin plots, for all cell types, after short stress exposure. D. Mitochondria metrics related to the Cristae Orientation of the mitochondria (Cristae Orientation Major, Cristae Orientation Minor, Cristae Orientation Angle, Cristae Area) are displayed using violin plots, for all cell types, after short stress exposure. See Table S1 for metrics description and y-axis units.

**Figure S7:**
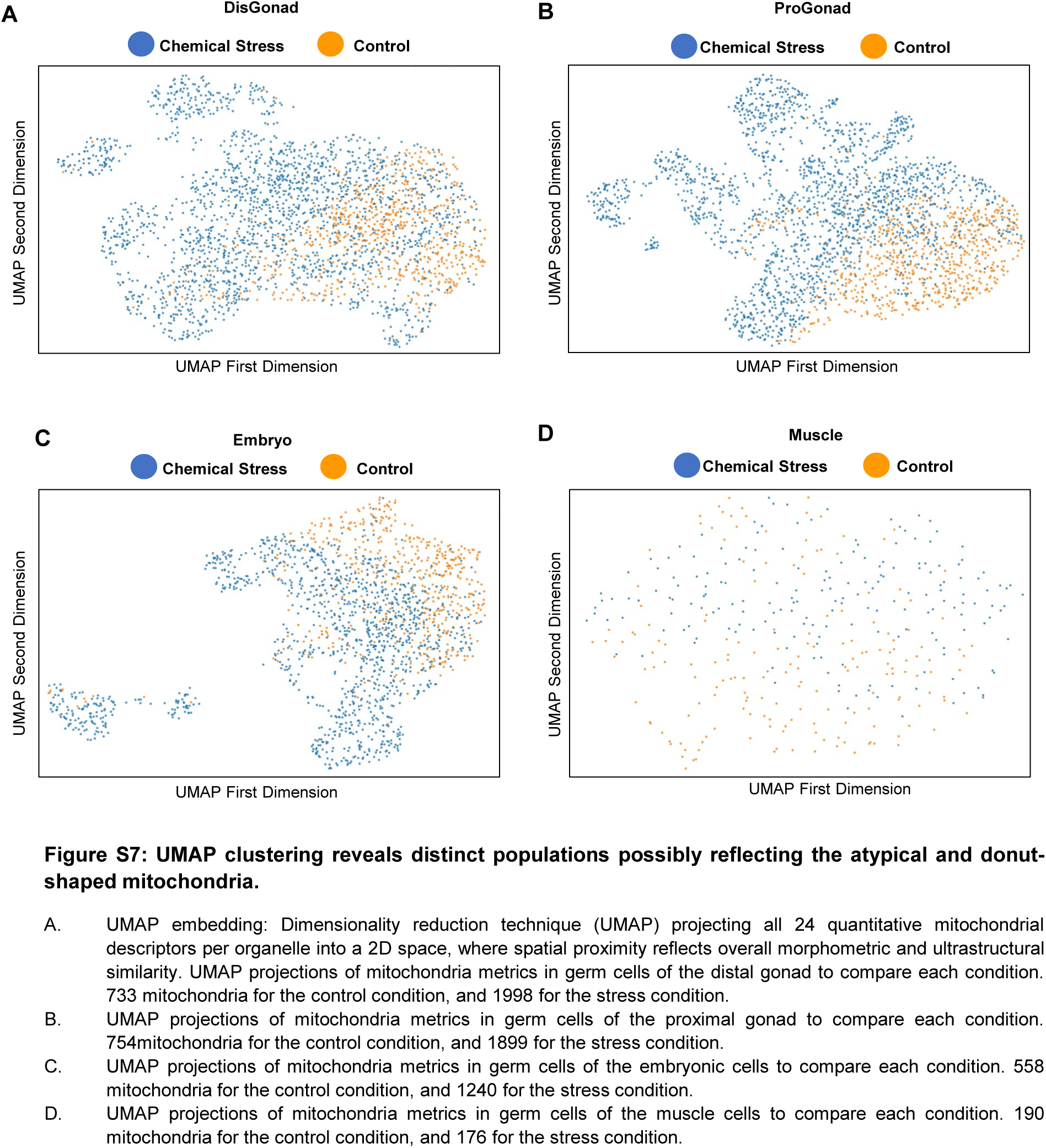
UMAP clustering reveals distinct populations possibly reflecting the atypical and donut-shaped mitochondria. A. UMAP embedding: Dimensionality reduction technique (UMAP) projecting all 24 quantitative mitochondrial descriptors per organelle into a 2D space, where spatial proximity reflects overall morphometric and ultrastructural similarity. UMAP projections of mitochondria metrics in germ cells of the distal gonad to compare each condition. 733 mitochondria for the control condition, and 1998 for the stress condition. B. UMAP projections of mitochondria metrics in germ cells of the proximal gonad to compare each condition. 754mitochondria for the control condition, and 1899 for the stress condition. C. UMAP projections of mitochondria metrics in germ cells of the embryonic cells to compare each condition. 558 mitochondria for the control condition, and 1240 for the stress condition. D. UMAP projections of mitochondria metrics in germ cells of the muscle cells to compare each condition. 190 mitochondria for the control condition, and 176 for the stress condition.

**Supplementary Table 1:**
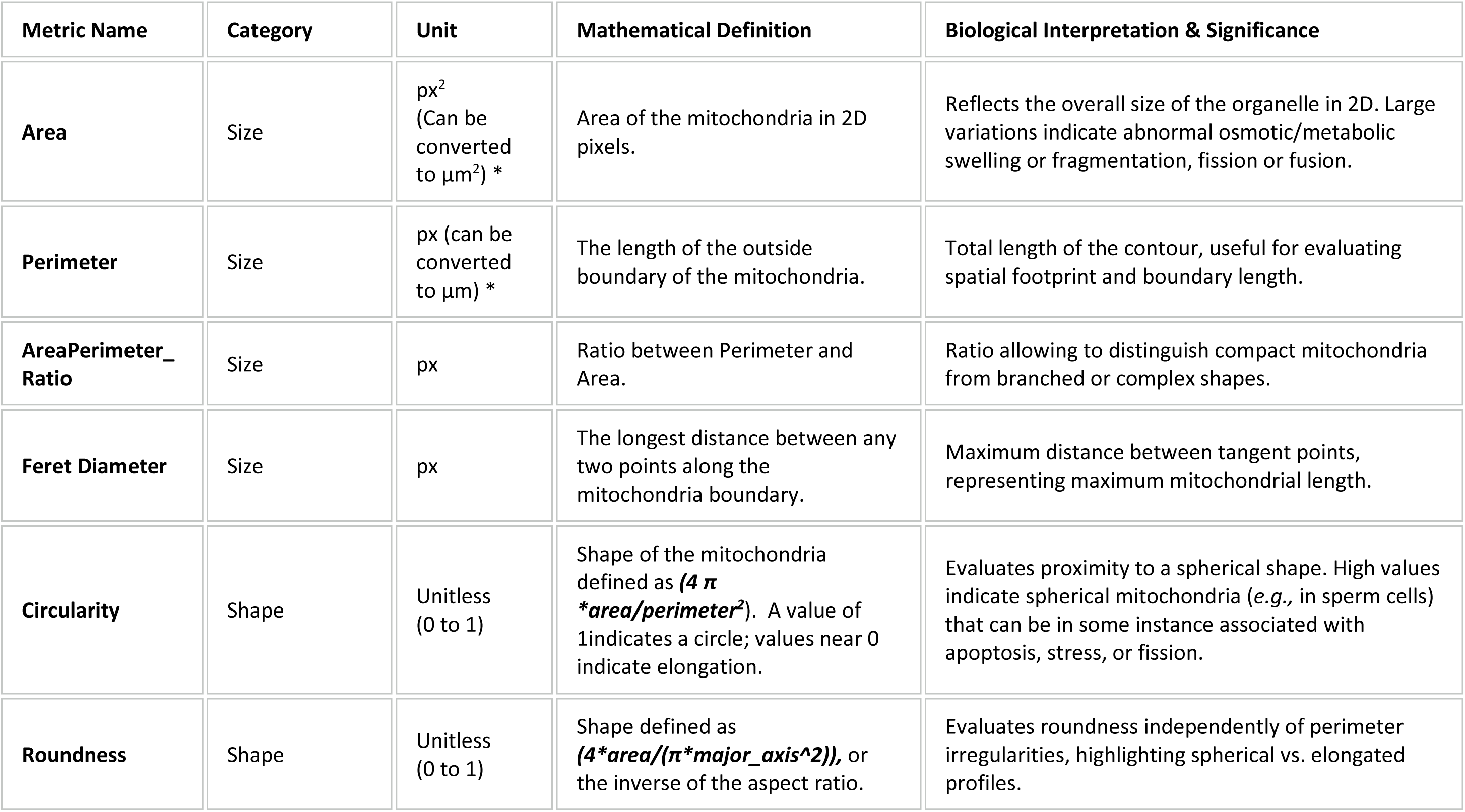
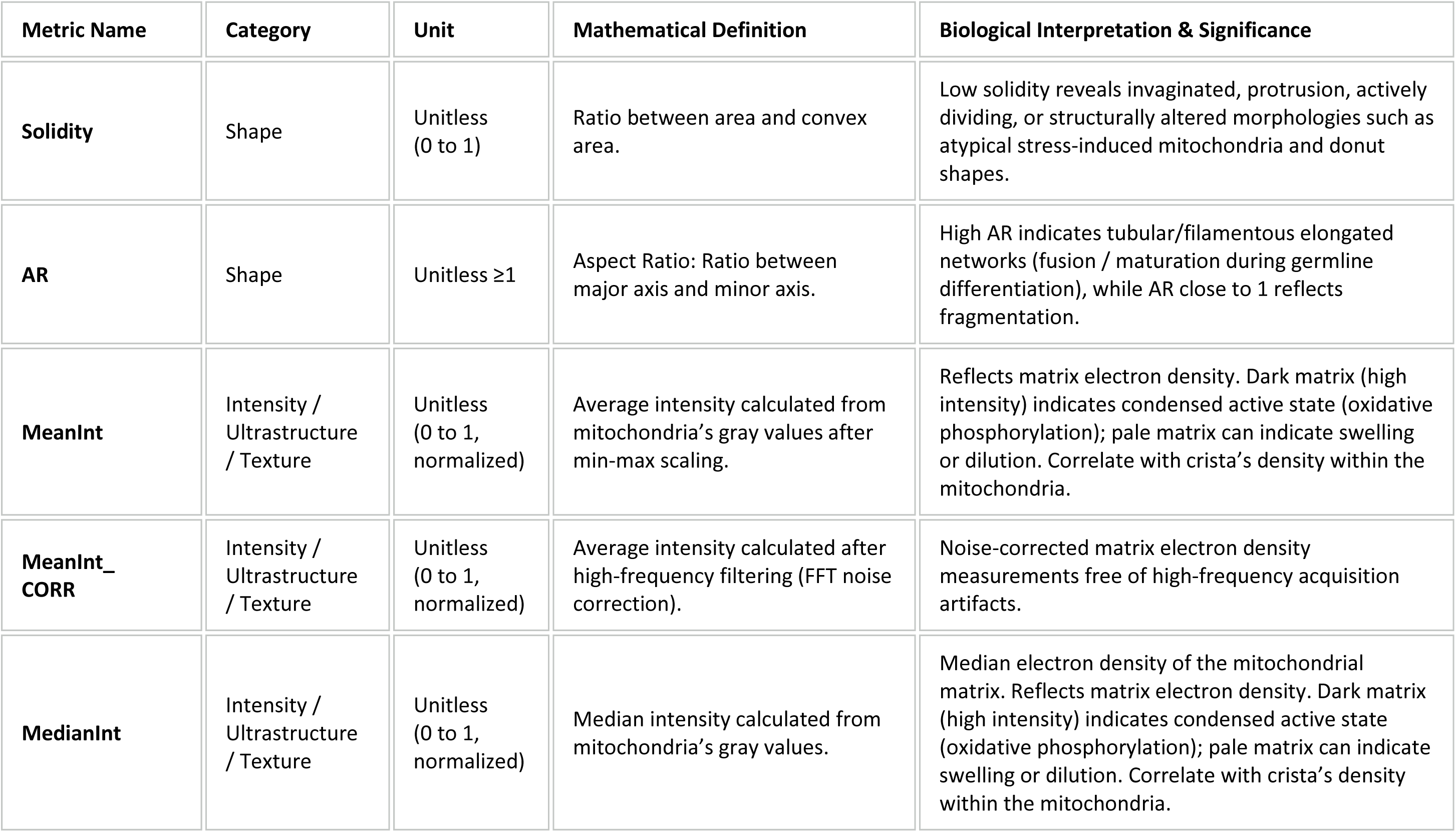
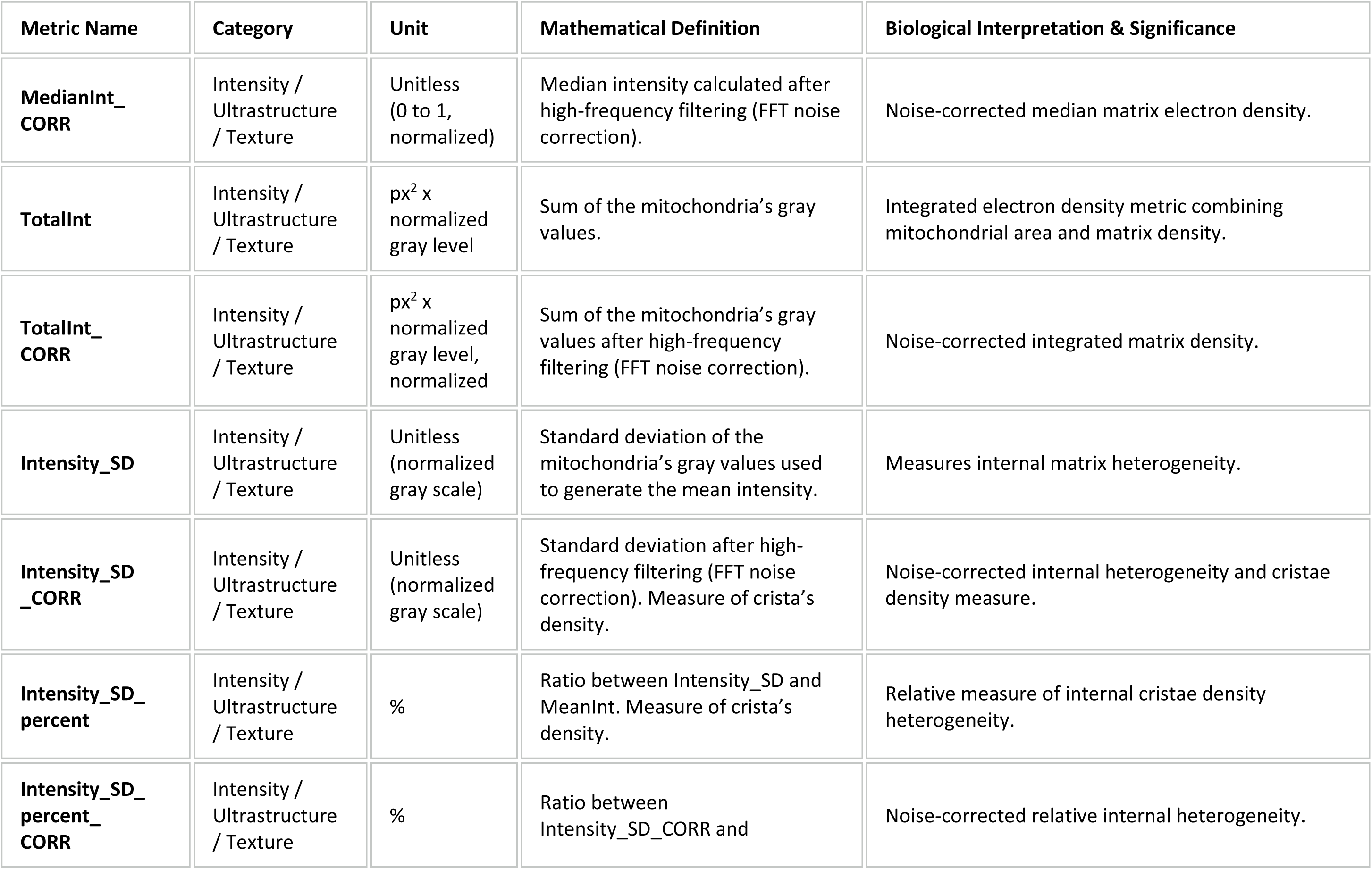
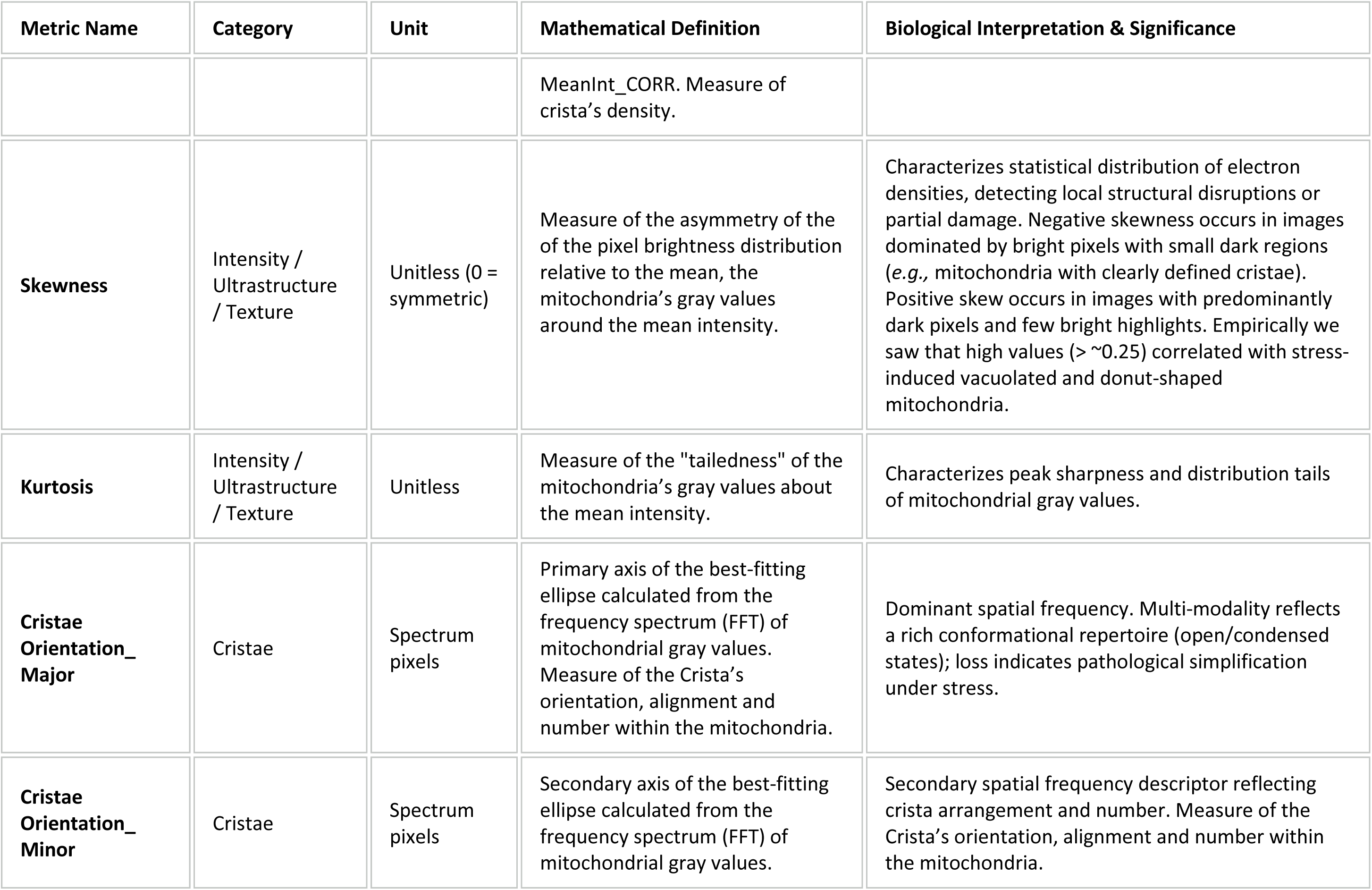
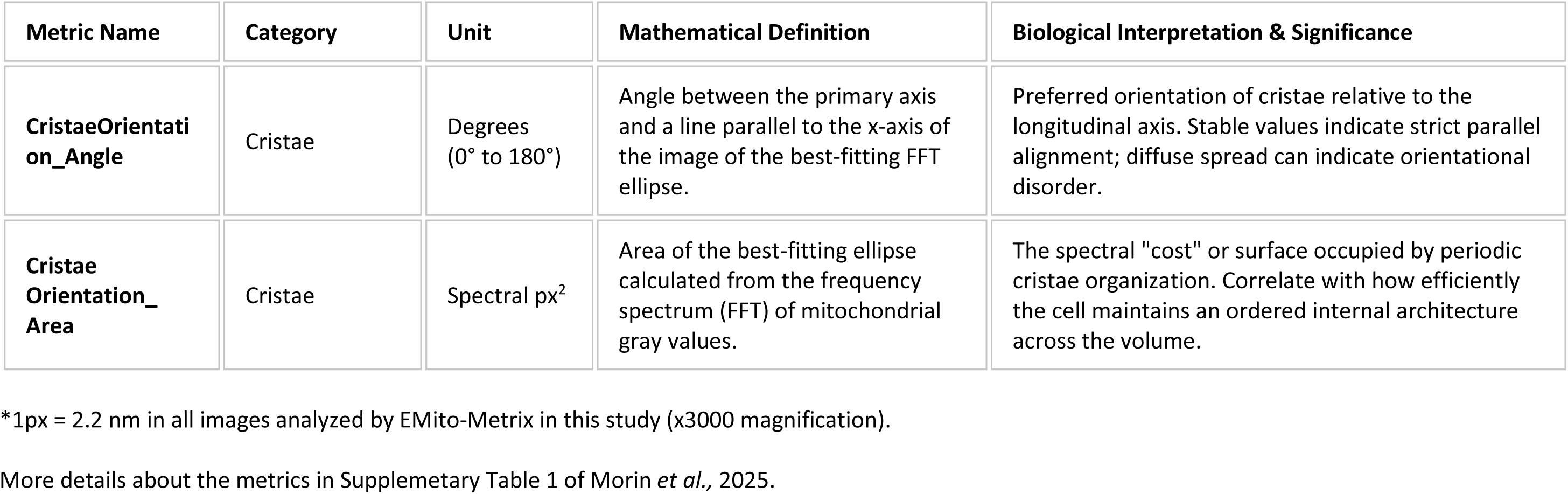
Glossary of EMito-Metrics names.

| Metric Name | Category | Unit | Mathematical Definition | Biological Interpretation & Significance |
|---|---|---|---|---|
| **Area** | Size | px<sup>2</sup><br>(Can be converted to µm<sup>2</sup>) * | Area of the mitochondria in 2D pixels. | Reflects the overall size of the organelle in 2D. Large variations indicate abnormal osmotic/metabolic swelling or fragmentation, fission or fusion. |
| **Perimeter** | Size | px (can be converted to µm) * | The length of the outside boundary of the mitochondria. | Total length of the contour, useful for evaluating spatial footprint and boundary length. |
| **AreaPerimeter_Ratio** | Size | px | Ratio between Perimeter and Area. | Ratio allowing to distinguish compact mitochondria from branched or complex shapes. |
| **Feret Diameter** | Size | px | The longest distance between any two points along the mitochondria boundary. | Maximum distance between tangent points, representing maximum mitochondrial length. |
| **Circularity** | Shape | Unitless<br>(0 to 1) | Shape of the mitochondria defined as ***(4 π \*area/perimeter<sup>2</sup>***). A value of 1indicates a circle; values near 0 indicate elongation. | Evaluates proximity to a spherical shape. High values indicate spherical mitochondria (*e.g.,* in sperm cells) that can be in some instance associated with apoptosis, stress, or fission. |
| **Roundness** | Shape | Unitless<br>(0 to 1) | Shape defined as ***(4\*area/(π\*major_axis^2)),*** or the inverse of the aspect ratio. | Evaluates roundness independently of perimeter irregularities, highlighting spherical vs. elongated profiles. |
| Metric Name | Category | Unit | Mathematical Definition | Biological Interpretation & Significance |
| **Solidity** | Shape | Unitless<br>(0 to 1) | Ratio between area and convex area. | Low solidity reveals invaginated, protrusion, actively dividing, or structurally altered morphologies such as atypical stress-induced mitochondria and donut shapes. |
| **AR** | Shape | Unitless ≥1 | Aspect Ratio: Ratio between major axis and minor axis. | High AR indicates tubular/filamentous elongated networks (fusion / maturation during germline differentiation), while AR close to 1 reflects fragmentation. |
| **MeanInt** | Intensity / Ultrastructure / Texture | Unitless<br>(0 to 1, normalized) | Average intensity calculated from mitochondria’s gray values after min-max scaling. | Reflects matrix electron density. Dark matrix (high intensity) indicates condensed active state (oxidative phosphorylation); pale matrix can indicate swelling or dilution. Correlate with crista’s density within the mitochondria. |
| **MeanInt_CORR** | Intensity / Ultrastructure / Texture | Unitless<br>(0 to 1, normalized) | Average intensity calculated after high-frequency filtering (FFT noise correction). | Noise-corrected matrix electron density measurements free of high-frequency acquisition artifacts. |
| **MedianInt** | Intensity / Ultrastructure / Texture | Unitless<br>(0 to 1, normalized) | Median intensity calculated from mitochondria’s gray values. | Median electron density of the mitochondrial matrix. Reflects matrix electron density. Dark matrix (high intensity) indicates condensed active state (oxidative phosphorylation); pale matrix can indicate swelling or dilution. Correlate with crista’s density within the mitochondria. |
| Metric Name | Category | Unit | Mathematical Definition | Biological Interpretation & Significance |
| **MedianInt_CORR** | Intensity / Ultrastructure / Texture | Unitless<br>(0 to 1, normalized) | Median intensity calculated after high-frequency filtering (FFT noise correction). | Noise-corrected median matrix electron density. |
| **TotalInt** | Intensity / Ultrastructure / Texture | px<sup>2</sup> x normalized gray level | Sum of the mitochondria’s gray values. | Integrated electron density metric combining mitochondrial area and matrix density. |
| **TotalInt_CORR** | Intensity / Ultrastructure / Texture | px<sup>2</sup> x normalized gray level, normalized | Sum of the mitochondria’s gray values after high-frequency filtering (FFT noise correction). | Noise-corrected integrated matrix density. |
| **Intensity_SD** | Intensity / Ultrastructure / Texture | Unitless (normalized gray scale) | Standard deviation of the mitochondria’s gray values used to generate the mean intensity. | Measures internal matrix heterogeneity. |
| **Intensity_SD_CORR** | Intensity / Ultrastructure / Texture | Unitless (normalized gray scale) | Standard deviation after high-frequency filtering (FFT noise correction). Measure of crista’s density. | Noise-corrected internal heterogeneity and cristae density measure. |
| **Intensity_SD_percent** | Intensity / Ultrastructure / Texture | % | Ratio between Intensity_SD and MeanInt. Measure of crista’s density. | Relative measure of internal cristae density heterogeneity. |
| **Intensity_SD_percent_CORR** | Intensity / Ultrastructure / Texture | % | Ratio between Intensity_SD_CORR and | Noise-corrected relative internal heterogeneity. |
| Metric Name | Category | Unit | Mathematical Definition | Biological Interpretation & Significance |
|  |  |  | MeanInt_CORR. Measure of crista’s density. |  |
| **Skewness** | Intensity / Ultrastructure / Texture | Unitless (0 = symmetric) | Measure of the asymmetry of the of the pixel brightness distribution relative to the mean, the mitochondria’s gray values around the mean intensity. | Characterizes statistical distribution of electron densities, detecting local structural disruptions or partial damage. Negative skewness occurs in images dominated by bright pixels with small dark regions (*e.g.,* mitochondria with clearly defined cristae). Positive skew occurs in images with predominantly dark pixels and few bright highlights. Empirically we saw that high values (> ~0.25) correlated with stress-induced vacuolated and donut-shaped mitochondria. |
| **Kurtosis** | Intensity / Ultrastructure / Texture | Unitless | Measure of the "tailedness" of the mitochondria’s gray values about the mean intensity. | Characterizes peak sharpness and distribution tails of mitochondrial gray values. |
| **Cristae Orientation_Major** | Cristae | Spectrum pixels | Primary axis of the best-fitting ellipse calculated from the frequency spectrum (FFT) of mitochondrial gray values. Measure of the Crista’s orientation, alignment and number within the mitochondria. | Dominant spatial frequency. Multi-modality reflects a rich conformational repertoire (open/condensed states); loss indicates pathological simplification under stress. |
| **Cristae Orientation_Minor** | Cristae | Spectrum pixels | Secondary axis of the best-fitting ellipse calculated from the frequency spectrum (FFT) of mitochondrial gray values. | Secondary spatial frequency descriptor reflecting crista arrangement and number. Measure of the Crista’s orientation, alignment and number within the mitochondria. |
| Metric Name | Category | Unit | Mathematical Definition | Biological Interpretation & Significance |
| **CristaeOrientation_Angle** | Cristae | Degrees<br>(0° to 180°) | Angle between the primary axis and a line parallel to the x-axis of the image of the best-fitting FFT ellipse. | Preferred orientation of cristae relative to the longitudinal axis. Stable values indicate strict parallel alignment; diffuse spread can indicate orientational disorder. |
| **Cristae Orientation_Area** | Cristae | Spectral px<sup>2</sup> | Area of the best-fitting ellipse calculated from the frequency spectrum (FFT) of mitochondrial gray values. | The spectral "cost" or surface occupied by periodic cristae organization. Correlate with how efficiently the cell maintains an ordered internal architecture across the volume. |

\*1px = 2.2 nm in all images analyzed by EMito-Metrix in this study (x3000 magnification).
More details about the metrics in Supplemetary Table 1 of Morin *et al.,* 2025.

